# Computational basis of hierarchical and counterfactual information processing

**DOI:** 10.1101/2024.01.30.578076

**Authors:** Mahdi Ramadan, Cheng Tang, Nicholas Watters, Mehrdad Jazayeri

## Abstract

Cognitive theories attribute humans’ unparalleled capacity in solving complex multistage decision problems to distinctive hierarchical and counterfactual reasoning strategies. Here, we used a combination of human psychophysics and behaviorally-constrained neural network modeling to understand the computational basis of these cognitive strategies. We first developed a multi-stage decision-making task that humans solve using a combination of hierarchical and counterfactual processing. We then used a series of hypothesis-driven behavioral experiments to systematically dissect the potential computational constraints that underlie these strategies. One experiment revealed that humans have limited capacity for parallel processing. Another indicated that counterfactuals do not fully compensate for this limitation because of working memory limits. A third experiment revealed that the degree to which humans use counterfactuals depends on the fidelity of their working memory. Next, we asked whether the strategies humans adopt are computationally rational; i.e., optimal under these constraints. To do so, we analyzed the behavior of a battery of task-optimized recurrent neural networks (RNNs) that were subjected to one or more of these constraints. Remarkably, only RNNs that were subjected to all these constraints behaved similarly to humans. Further analysis of the RNNs revealed that what cognitive theories posit as distinctive strategies such as hierarchical and counterfactual are subdivisions in a continuum of computationally rational solutions that includes optimal, counterfactual, postdictive, and hierarchical.

## Introduction

The human brain finds solutions to complex multistage decision problems that are far more flexible than those learned by artificial systems. Cognitive theories attribute this flexibility to specific algorithms such as hierarchical information processing and counterfactual reasoning (Huys et al., 2015; Lake et al., 2017; Tenenbaum et al., 2011; Zylberberg, 2021). Counterfactual reasoning is an important building block of our mental landscape enabling us to imagine alternative accounts of our prior experiences (Gerstenberg et al., 2020; Pearl, 2009; Van Hoeck et al., 2015). A familiar scenario is when we feel the need to revisit our assumptions due to an unexpected turn of events, for example, an unexpected fork in the road while driving, an abrupt emotional reaction from a friend, or a twist in the storyline of a book or movie. These situations compel us to review past events and assumptions and look for alternative interpretations that could provide a plausible explanation for unexpected observations. Given the centrality of counterfactuals in human cognition, it behooves us to understand the computational underpinnings of this strategy.

Humans commonly rely on counterfactuals when facing decision trees with a hierarchy of if-then scenarios leading to different outcomes (Buchsbaum et al., 2012; Gerstenberg, 2022; Lucas & Kemp, 2015; Sloman & Hagmayer, 2006; Zylberberg et al., 2018). However, what is puzzling is that decision trees do not inherently require computing counterfactuals. On the contrary, the optimal strategy for solving a decision tree is to use each observation to update the posterior belief over all states within the tree (Kuperwajs & Ma, 2021; Purcell & Kiani, 2016; Zheng et al., 2022). However, for sufficiently complex decision trees, an optimal posterior updating strategy can become extremely demanding and sometimes intractable (Gershman et al., 2015; Van Opheusden et al., 2017). Curiously, these are also the conditions for which humans typically use counterfactuals. These considerations raise several interrelated questions about counterfactual reasoning. Is this an *ex-post* strategy that humans use as a crutch for updating the posterior belief in complex decision trees? Do counterfactuals introduce sub-optimalities in behavior? If so, what are they, and do humans adopt a computationally rational approach to use counterfactuals effectively?

To answer these questions we developed a moderately challenging multi-stage decision-making task for evaluating the computational basis of cognitive strategies in humans. Briefly, participants had to infer the position of an invisible moving ball within an H-shaped maze using partial and uncertain cues about its position (Fig. 1a). Comparing participants’ behavior to inference models implementing various cognitive algorithms, we found that humans use a hierarchical strategy to solve the task sequentially, and when uncertain, revise their decisions by considering counterfactuals. Additionally, participants’ reaction time profiles and eye movements provided evidence that they relied on counterfactuals to revise their decisions.

**Figure 1.**
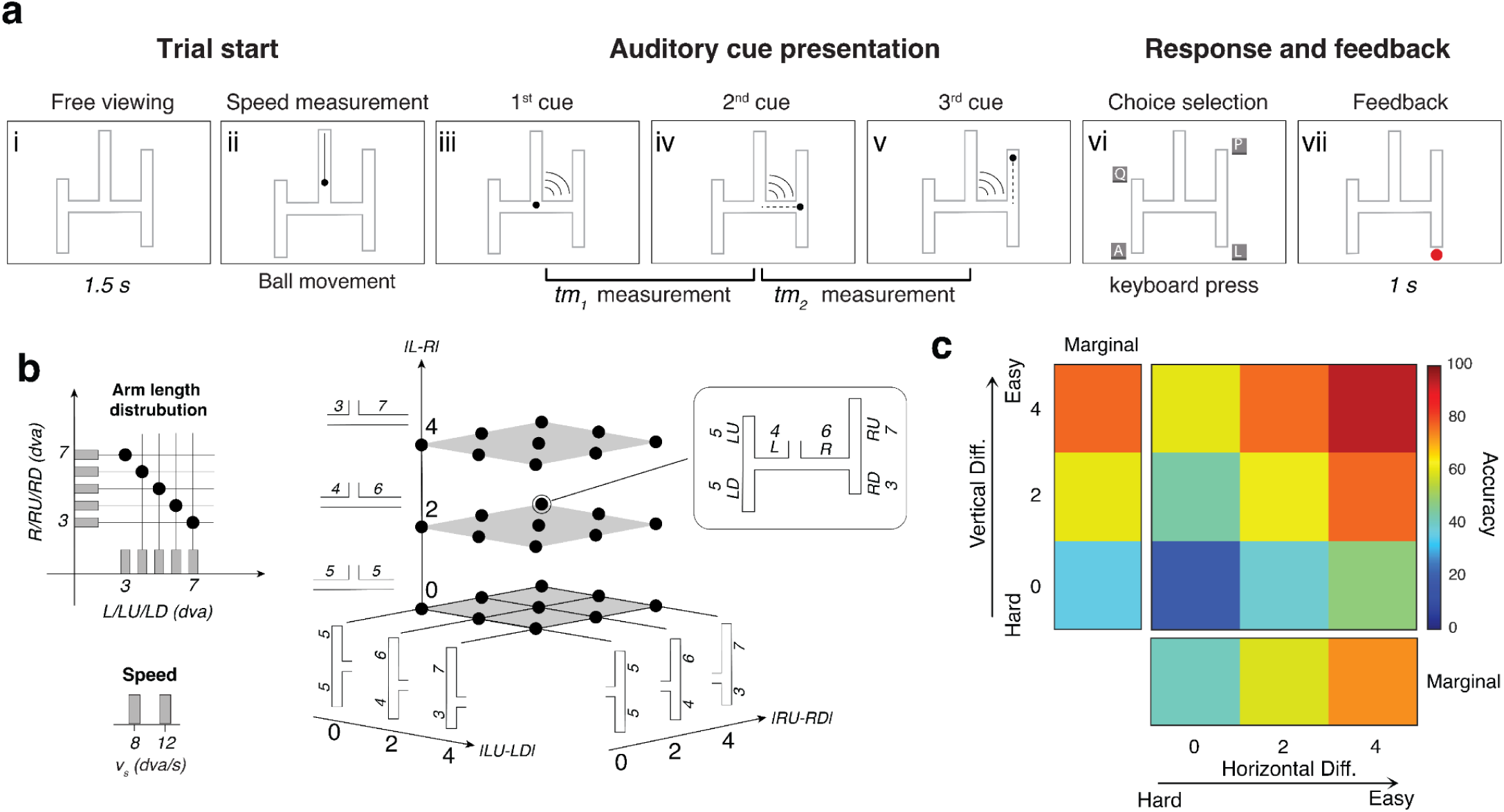
The H-Maze task and performance. **(a)** Task. (i) Participant is presented with an H-shaped maze and four possible choices. (ii) A ball moves visibly toward the maze with constant speed. (iii) An auditory cue signals when the ball reaches the horizontal segment at which time the ball is made invisible and continues to move leftward or rightward. (iv) A second auditory cue signals when the ball reaches the vertical segment and the ball changes direction upward or downward (*tm_1_*: the time between the first and second cues). (v) A third auditory cue signaled when the ball reached the endpoint (*tm_2_*: the time between the second and third cues). (vi) Participant presses one of four keyboard buttons to indicate their belief about the final exit point. (vii) Participant receives binary visual feedback only at the location of their choice. **(b)** H-maze arm lengths, denoted by *L* (left)*, R* (right)*, LU* (left-up)*, LD* (left-down)*, RU* (right-up), and *RD* (right-down), were sampled from a discrete uniform distribution with values 3, 4, 5, 6, and 7 degree visual angle (dva) subject to the constraint that the sum of all arm pairs (*L+R, LU+LD, RU+RD*) was 10 dva (top left). Accordingly, the absolute difference between each arm pair (|*L - R|,* |*LU - LD|,* |*RU - RD|*) could be either 0 (5, 5), 2 (6, 4), or 4 (7, 3) dva (right). The H-maze in the box shows an example H-maze subjected to these constraints. The speed of the ball was either 8 or 12 dva/s, chosen randomly (bottom left). **(c)** Performance averaged across participants. Percentage of correct responses as a function of absolute length difference between the two horizontal arms (|L-R|) and the two vertical arms associated with the correct horizontal arm (|LU-RD| for left and |RU-RD| for right).

Next, we used a set of experiments to reverse engineer the computational constraints from which these algorithms derive. Experiment 1 indicated that the hierarchical strategy results from a bottleneck associated with processing parallel streams of evidence. Experiment 2 indicated that compensatory counterfactuals are imperfect due to working memory limits. Experiment 3 indicated that humans are computationally rational in that the degree to which they rely on counterfactuals depends on the fidelity of their working memory.

Next, we used a modeling approach to test the importance of these computational constraints on adopting a counterfactual strategy. We trained multiple artificial recurrent neural network (RNN) models to perform the same task as humans and subjected each to different subsets of those constraints. Unconstrained models adopted an optimal strategy that deviated significantly from the human participants’ behavior. The addition of constraints altered the model’s behavior. For example, a processing bottleneck constraint shifted the model’s responses toward a hierarchical strategy and working memory limits impact the degree to which the model relied on counterfactuals. Remarkably, the model that most accurately emulated humans’ behavioral responses patterns was the one that was subjected to all the constraints inferred from human’s behavior. Finally, parametric analysis of the models revealed that distinct cognitive algorithms such as optimal, counterfactual, postdictive, and hierarchical may be more accurately characterized as subdivisions in a continuum that neural systems may adopt depending on the task and computational constraints.

## Results

We report the results of our experiments in the order that they were performed. We first tested a small sample of participants to establish our hypotheses, then pre-registered our hypotheses and analyses in the Open Science Framework (*OSF*, 2024.), and then repeated all experiments with pre-registered analyses in a large cohort of participants recruited online.

### The H-Maze Task

On each trial, a ball approaches the horizontal segment of an H-shaped maze from above. Upon reaching the maze, the ball becomes invisible and moves along either the left or right horizontal arm. Upon reaching the corresponding vertical segment, the ball turns upward or downward along one of the vertical arms, and stops after reaching the corresponding exit. After the ball stops, participants must report the exit point. Participants must infer the ball’s trajectory using partial and ambiguous information provided by three brief auditory clicks. The first click occurs when the ball turns into one of the horizontal arms, the second, when it enters one of the vertical arms, and the third, when it reaches the exit. The time between the first and second clicks (*t_h_*) provides information about the horizontal arm the ball enters, and the time between the second and third clicks (*t_v_*), provides information about the subsequent vertical arm. Participants must measure these two time intervals and choose the path whose horizontal and vertical arms were compatible with those measurements. In summary, participants have to infer the correct path from the relative timing of the auditory clicks. Crucially, we varied the lengths of both horizontal and vertical arms on a trial-by-trial basis making it essential for participants to use a flexible dynamic inference strategy (Fig. 1b). Throughout the manuscript, we will denote the length of the horizontal arms by *L* (left arm) and *R* (right arm), and the four vertical arms by *LU* (left-up)*, LD* (left-down)*, RU* (right-up)*, RD* (right-down). We may also express arm lengths in units of time reflecting how long it would take for the moving ball to traverse that arm. When doing so, we will use the corresponding labels *t_L_* (left arm), *t_R_*(right arm), *t_LU_* (left-up), *t_LD_* (left-down), *t_RU_* (right-up), and *t_RD_* (right-down).

Analysis of behavior indicated that participants learned the task and were able to use the timing cues to infer the exit point. Average performance across participants improved as a function of the difference between both the horizontal and vertical arms (Fig. 1c). We therefore proceeded with a model-based analysis of behavior to infer the participant’s cognitive strategy.

### Models of different cognitive computational strategies

To quantitatively investigate the strategy participants used for solving the task, we developed a range of models implementing different cognitive strategies with different degrees of optimality. Our first model, which we refer to as the “optimal” model, chooses the exit point based using a maximum-likelihood strategy; it chooses the path whose composition of horizontal and vertical arms is most consistent with the joint distribution of *t_h_* and *t_v_* (Fig. 2a, Top-Left). Our second model implements a hierarchical strategy by breaking down the problem into two hierarchically organized sequential decisions (Fig. 2a, Bottom-Left). It first uses *t_h_* to choose between the horizontal arms, and then uses *t_v_* to choose between the corresponding vertical arms. This strategy is suboptimal because it does not allow the decision about the first arm to benefit from information about the second arm and vice versa. For example, the model may commit to the left horizontal arm based on *t_h_*even if *t_v_* is not consistent with either of the two left vertical arms. Our third model implements a postdictive inference strategy (Fig. 2a, Top-Right). This model functions hierarchically but postpones its first decision until after it has received information about the second arm. More specifically, the model’s decision about the horizontal arm is conditioned not only on *t_h_* but also on the degree to which *t_v_* is consistent with the corresponding vertical arms. Note that this differs from the optimal model because the postdictive model makes a horizontal arm choice based on the sum of the likelihoods of the two endpoints for each horizontal arm, hence may be suboptimal when the maximum-likelihood endpoint is on the same horizontal branch as the minimum-likelihood endpoint. Fourth, we developed a hierarchical model that would additionally consider counterfactuals (Fig. 2a, Bottom-Right). This model starts with the two-stage hierarchical strategy. However, if the likelihood of the vertical arms in the second stage is below a certain threshold, it revises its first decision and chooses between the other two vertical arms. In general, counterfactual reasoning is suboptimal because recalling information from memory is imperfect (Ratcliff, 1978; Shadlen & Shohamy, 2016). Note that revisions in counterfactual models differ qualitatively from vacillations that occur during simple decision-making (Peixoto et al., 2021; Resulaj et al., 2009). Vacillations arise from fluctuation of evidence, whereas counterfactuals arise from a deliberate process that uses late evidence to revise earlier commitments.

**Figure 2.**
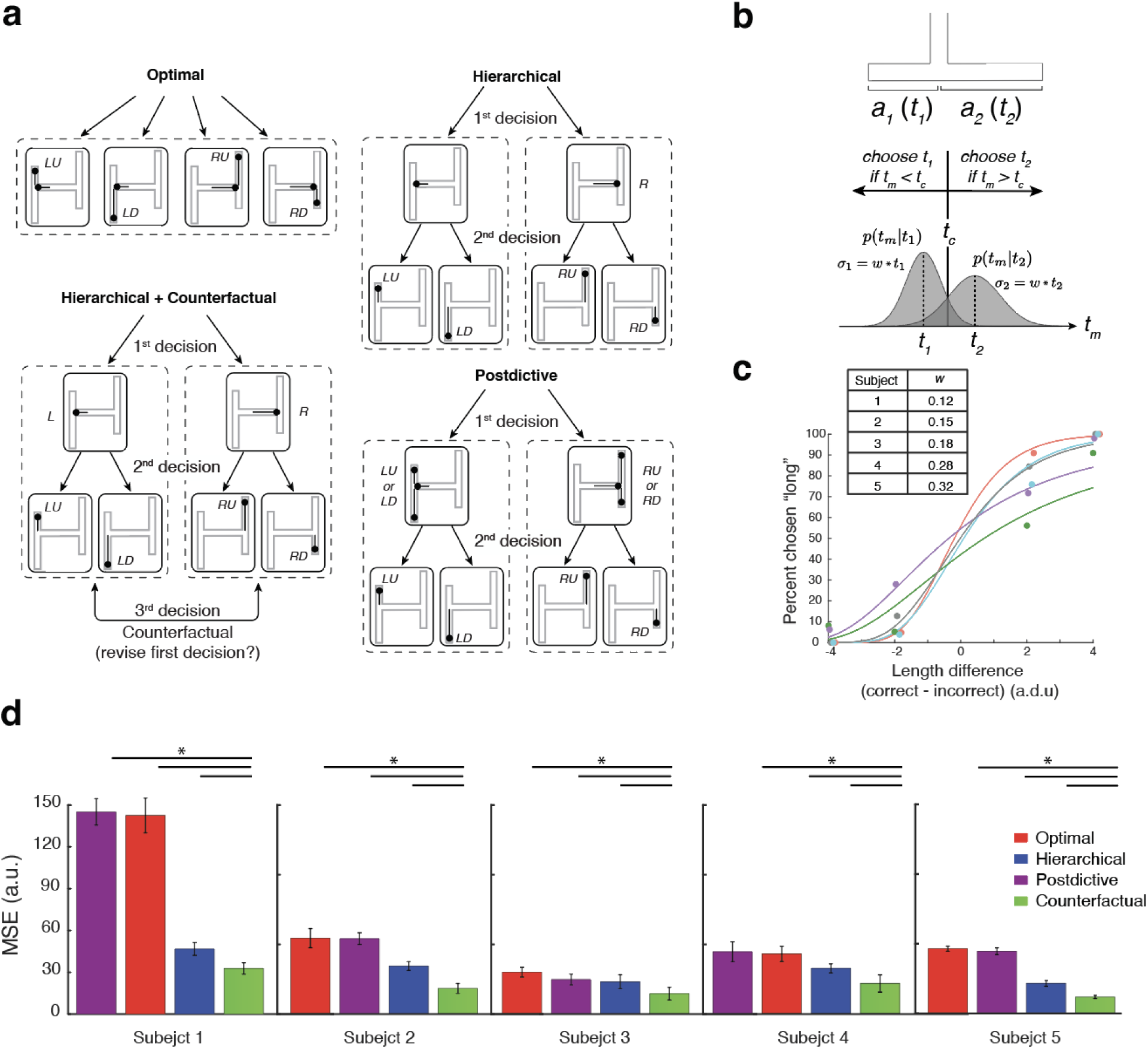
Model-based analysis of behavior. **(a)** Schematics showing models implementing different computational strategies to solve the H-maze task. Arrows indicate potential decisions and dashed boxes enclose the hypotheses being considered at each stage. The black lines with a ball attached (inside H-maze) indicate which arm segments are being evaluated. Top-Left: The optimal model, which infers the exit based on the joint likelihood of two time intervals measured between consecutive clicks. Top-Right: The hierarchical model, which infers the exit by first choosing between the two horizontal arms using the first and second clicks, and then between the corresponding vertical arms using the second and third clicks. Bottom-Right: The postdictive model, which behaves similarly to the hierarchical model with the key difference that it chooses the horizontal arm postdictively using both time intervals. Bottom-Left: The counterfactual model. This model behaves similarly to the hierarchical model but when the likelihoods for both vertical arms under consideration are below a threshold, the model reevaluates the evidence in the context of the alternative horizontal arm, which may lead to a revision of the decision about the horizontal arm. **(b)** The decision model. Left: The ball takes one of two arms with lengths *a_1_* and *a_2_*. Since the ball moves at a constant speed, the two arm lengths correspond to two specific time intervals, *t_1_* and *t_2_*. Right: The participant makes a noisy measurement, *t_m,_* and uses the two conditional probabilities, *p(t_m_|t_1_)* and *p(t_m_|t_2_),* to choose between the two arms. We modeled the measurement noise as a zero-mean Gaussian distribution whose standard deviation scales with the base interval (*t_1_* and *t_2_*), with the constant of proportionality *w* (*σ=wt*). The two distributions cross over at the criterion, *t_c_*. The model chooses *t_1_* when *t_m_* is smaller than *t_c_* and *t_2_*, otherwise. We used this model to measure each animal’s timing variability (weber fraction, *w*). **(c)** Performance and model fit for the T-maze task. Participants’ performance is plotted in terms of the percent of trials the longer arm was chosen as a function of the length difference between the correct and incorrect arms (colored circles). Model fits for each participant are shown as lines. Table: fitted *w* values for each of the five participants. **(d)** The mean squared error computed between models’ and participants’ choice probability (see choice analysis in Methods), cross-validated five-fold (errorbars: SEM). Pairwise comparisons (lines on the top) indicated that the MSE of the counterfactual model was significantly lower than other models in capturing participants’ choice patterns (Left-Tailed T-test, p<0.05).

### Humans use hierarchical reasoning and counterfactuals to solve the task

Next we compare participants’ performance against the models across different H-Maze geometries. If we assume that models can measure *t_h_*and *t_v_* perfectly, their performance would be trivially at 100%. To make the models’ more comparable to humans, we assumed that time measurements are noisy. Following the scalar property of timing in humans (Malapani & Fairhurst, 2002), we modeled noise in timing with a Gaussian distribution whose standard deviation scales with the base interval with a scale factor known as the weber fraction, *w*. We fit *w* for each participant on an independent T-Maze control task (Fig. 2b, Top) in which participants had to discriminate between two horizontal arms associated with two time intervals (*t_1_* and *t_2_*). Under the assumption of scalar Gaussian noise (Fig. 2b, Middle), we computed the maximum likelihood estimate of *w* based on the participant’s performance in the T-maze (Fig. 2b, Bottom). We used the fitted *w* values for each participant (Fig. 2c, inset) for all subsequent model comparisons. Note that we estimated *w* for each participant using an independent measure based on the performance in the T-maze task. This approach enabled us to compare participants’ behavior in the H-maze task to cognitive models (optimal, hierarchical, etc.) without the need to additionally estimate each participant’s timing accuracy, providing a more rigorous approach to model comparison.

To compare the behavior of the models and participants, we proceeded with a more granular analysis that moved beyond a simple accuracy measure, and instead, examined the full pattern of participants’ choices (see Methods). We encoded response patterns in terms of four probabilities, one for each possible exit, denoted by *p_LU_*, *p_LD_*, *p_RU_*, and *p_RD_*. We measured these probabilities for participants and models across subsets of mazes with the same set of arm length differences and measured the Euclidean distance for choice patterns between models and participants. Finally, we compared the average mean squared error (MSE) between participants and models across all geometries. Results of this comparison indicated that the participants’ behavior was most accurately captured by the counterfactual model; that is, the full pattern of participants’ choices was most similar to that of the counterfactual model. For all participants, the counterfactual model had the smallest cross-validated MSE compared to the other models (Fig. 2d).

Note that with perfect memory, the counterfactual model behaves identically to the optimal model with the only difference that it evaluates the two sides of the maze sequentially. In reality, counterfactual reasoning may deviate from optimality because of the capacity limitations of working memory needed to recall and process past evidence. Accordingly, our counterfactual model had an additional parameter associated with working memory noise to account for how well participants recall past information. To account for the additional complexity of the counterfactual model, we performed all model comparisons using five-fold cross-validation.

We performed additional analyses to further test whether participants computed counterfactuals. One prominent feature of the counterfactual model is the presence of a third decision process associated with deciding whether to run a counterfactual, and if so subsequently evaluating the alternative arm segments before making a final choice (Fig. 2a). Since each decision increases processing time, an analysis of reaction times could provide additional evidence as to whether and when participants used a counterfactual strategy. An analysis of the counterfactual model indicated that counterfactuals are relied upon more often when the first decision is more difficult; i.e., the difference between the two horizontal arms is small. Accordingly, the hypothesis that participants compute counterfactuals predicts that reaction times would increase systematically with the degree of difficulty of the first decision. Consistent with this prediction, participants’ average reaction times increased in a graded fashion with the difficulty of the first decision (Supplementary Fig. 1). This finding provides further evidence that, when uncertain, participants computed counterfactuals.

We also analyzed participants’ eye movements to examine the strategy they used to perform the task. Previous studies have shown that the pattern of eye movements during spatial reasoning tasks can reveal behavioral strategy (Gerstenberg et al., 2017; Hayhoe & Ballard, 2005; Levy et al., 2009; Li et al., 2023; Liversedge & Findlay, 2000; Spering, 2022). Accordingly, we reasoned that shifts in gaze position over the H-maze would serve as another clue in deciphering participants’ strategy. Specifically, shifts of the eye toward the left or right early in the trial (e.g., after the 1st time interval) would serve as evidence for a hierarchical strategy and subsequent revisions of gaze position from one side to the other would serve as evidence for considering counterfactuals. Eye movements showed evidence for both hierarchical and counterfactual processes (Supplementary Fig. 2). Notably, the frequency of counterfactual eye movements was highest in trials with a difficult 1st decision in which computing counterfactuals was expected to occur most frequently (Supplementary Fig 3). Together, the model-based analysis of choices, the pattern of reaction times across conditions, and gaze positions provide strong converging evidence that participants’ strategy for solving the H-maze task is consistent with a combination of hierarchical and counterfactual decisions.

### Computational constraints on decision-making

Participants’ decision strategy raises important questions about the underlying computational constraints. First, why do participants not process the alternatives in parallel? Second, why are the counterfactual decisions suboptimal? Finally, is the participants’ reliance on suboptimal counterfactuals rational (i.e., do participants know when to rely on counterfactuals)? To address these questions, we conducted three additional experiments to address these questions and elucidate the constraints that underlie the participants’ strategy.

#### Task Variant 1

We reasoned that hierarchical information processing may be due to an inability to process multiple streams of evidence in parallel, in line with previous findings (Davis & Marcus, 2016; Ludwin-Peery et al., 2021; Smith et al., n.d.; Van Opheusden et al., 2017). To test this possibility, we developed a variant of the task that could only be solved by parallel processing. Participants were presented with four separate arms serving as conduits for four separate balls (Fig. 3a). Upon presentation of an auditory cue, the balls entered the four arms moving at constant speed. If a ball reached an end, it reversed direction and continued to move at the same speed. All balls kept moving until the presentation of a second auditory cue that coincided with one randomly chosen ball reaching an arm’s end. At that time, participants were offered four choices associated with the four arms and had to report the arm in which the ball was at an exit. Importantly, all task parameters including ball speed, arm lengths, and movement times were identical to the H-maze. Moreover, the introduction of direction reversion dissociated the balls’ position from time and forced a strategy in which all balls had to be tracked simultaneously, replicating the computational demands of solving the H-maze task using an optimal decision strategy. Participants’ performance on this task was slightly higher than chance level, but significantly lower than the performance of an ideal observer model solving experiment 1 using fitted *w* values measured for each participant separately on a control task (Fig. 3a, right; see T-Maze experiment in methods). The relatively low performance associated with parallel processing suggests that participants’ choice of a hierarchical strategy is due to a bottleneck associated with processing parallel streams of evidence.

**Figure 3.**
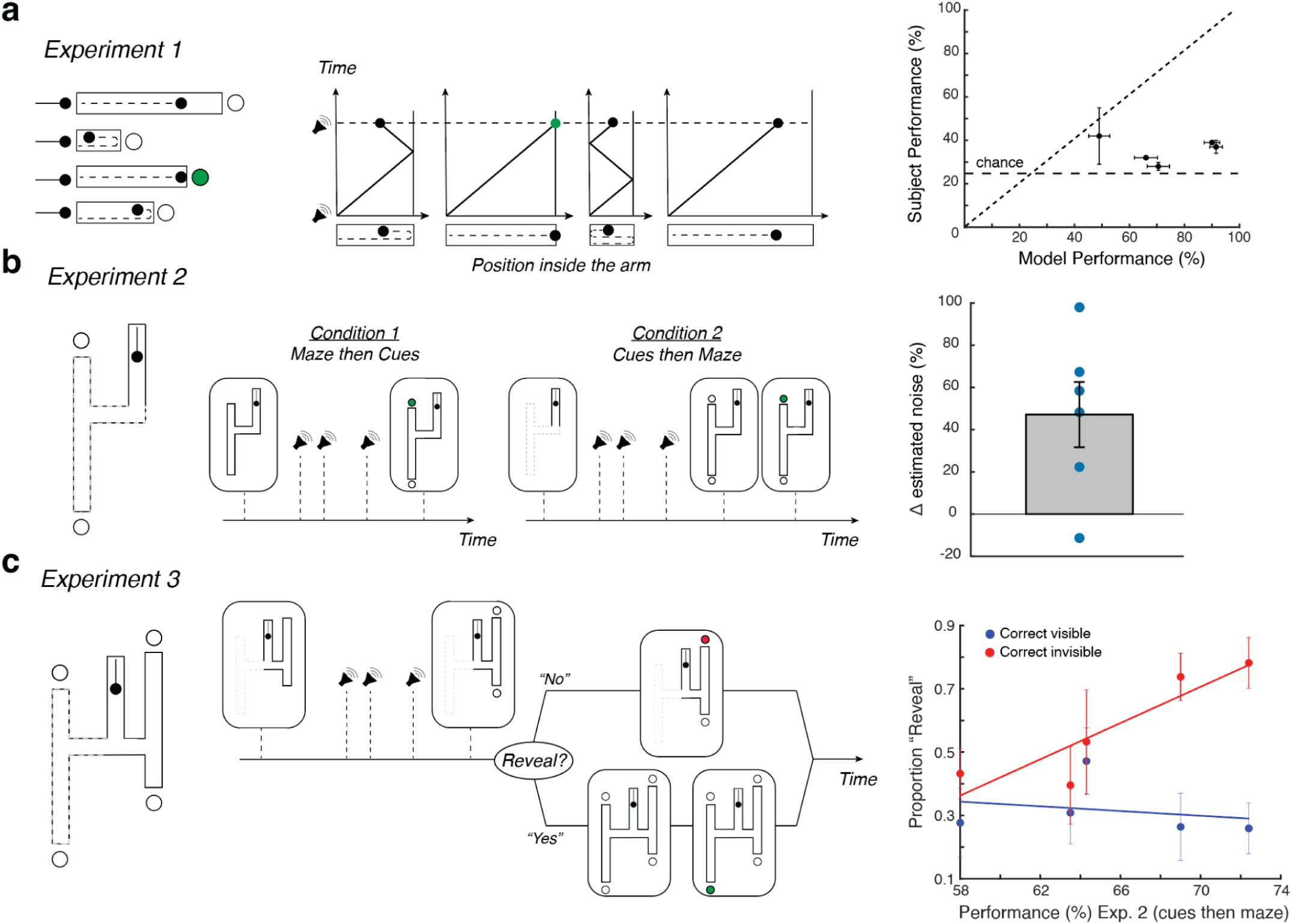
Task variants. **(a)** Task Variant 1. (Left) A variation in H-maze with independent arms. four balls moving at constant and identical speed enter four independent arms simultaneously. Upon entry, an auditory cue is played, and the balls become invisible. The balls continue to move inside their respective arm and reverse direction every time they reach an end. A second auditory cue is played when one of the balls, chosen at random, is at one end of its respective arm (left or right). Subsequently, participants are presented with four options and must choose the arm in which the ball is at its endpoint. Participants receive a binary visual feedback at the location of their choice. (Middle) Balls move back and forth inside their respective arms until the time of the second auditory cue. In the example shown, the correct answer is the second arm from the bottom. (Right) Performance for individual participants compared to an ideal observer model solving experiment 1 using fitted *w* values measured separately for each participant on a control task (horizontal black dashed line: chance-level performance; diagonal black dashed line: unity; vertical error bars: participant S.E.M over sessions; horizontal error bars: model S.E.M). Performance was below the level expected by an ideal observer (Left-Tailed T-Test: p<0.01 for all subjects, n=5). **(b)** Task Variant 2. (Left) A variation of H-maze with only the left half. (Middle) The experiment includes two conditions. In condition 1, the time course of the trial is identical to the original H-maze: stimulus, then the three auditory cues, followed by decision and feedback. In condition 2, participants first hear the three cues and then see the left half of the H-maze, followed by decision and feedback. (Right) The percent change in the estimated noise between condition 2 and condition 1. The estimated noise significantly increases in condition 2 compared to condition 1 (Blue circles: individual subjects, gray bar: mean percent change in estimated noise, error-bar: S.E.M across subjects, two-tailed T-Test: p = 0.028, n=6). **(c)** Task Variant 3. (Left) A variation on H-maze in which half of the H-maze is initially visible during the presentation of the auditory cues, and participants may choose to reveal the other half afterward. (Middle) After the presentation of one-half of the H-maze (left or right, chosen randomly) and the three auditory cues, participants may either choose between the two available exit points or ask to have the other half revealed and subsequently choose among the four exit points. (Right) The proportion of trials that participants asked for the other half of the H-maze to be revealed as a function of performance in Condition 2 of Experiment 2 (Cues then maze) across participants. Blue circles are trials where the correct answer was on the revealed half of the H-maze, and red circles are trials where the correct answer was on the unrevealed half of the H-maze. Error bars indicate s.d. across sessions, and the solid red and blue lines are the best linear fit to the data (Blue: r^2 = 0.05, p =0.70, Red: r^2 = 0.80, p = 0.039, n=5).

#### Task Variant 2

An ideal observer that relies on counterfactuals should, in principle, be able to attain optimal performance since counterfactuals can be used to evaluate all possibilities sequentially. However, to attain optimal performance, counterfactual processing should be noise-free, or else the performance would drop. Our analysis of participants’ behavior indicated that they relied on counterfactuals, but their performance was suboptimal, which led us to hypothesize that computing counterfactuals may incur additional noise. To test this hypothesis directly, we developed a variant of the task using a simpler T-maze geometry with two conditions (Fig. 3b). Condition 1 is analogous to the original H-maze task: the ball enters the maze through a vertical hallway and moves invisibly through the maze while auditory cues indicate the turning points and the time the ball reaches the one of the two exits, and participants are asked to report the exit point. In condition 2, we reversed the order of stimulus presentations such that the T-maze was presented *after* the presentation of the auditory cues. Importantly, in condition 2, participants must solve the task using counterfactuals; i.e., review past evidence from the auditory cues to evaluate alternatives offered by the subsequent presentation of the T-maze. The hypothesis that counterfactual processing is noisy predicts a higher estimated noise in condition 2 compared to condition 1. To test this, we took a model-based approach and computed the maximum likelihood estimate of each subject’s noise in both conditions based on the participant’s performance (see methods). Results were consistent with this prediction (Fig. 3b, right). Across participants, there was a significant increase in the estimated noise in condition 2 compared to condition 1 (two-tailed T-Test: p = 0.028, n=6), revealing the impact of counterfactual processing noise on performance.

#### Task Variant 3

Next, we examined the bounded rationality of participants while using counterfactuals. According to the bounded rationality hypothesis, participants should take into account the degrading effect of the counterfactual processing noise and titrate their reliance on counterfactuals accordingly. Specifically, this hypothesis predicts more reliance on counterfactuals for lower counterfactual processing noise and vice versa. To test this hypothesis, we leveraged the performance variance across participants in experiment 2, which varied widely between ∼58% to ∼74%. We asked the same participants to perform a variant of the H-maze task in which they could choose whether and when to rely on counterfactuals (Fig. 3c). In this variant, participants were presented with one side of the H-maze (left or right) during the presentation of the auditory cues. Afterward, participants were given an option to choose between two decision paths. They could either choose one of the two visible exit points as their final answer or ask for the other half of the H-maze to be revealed so that they can counterfactually evaluate the other exit points.

Remarkably, on trials where the correct exit point was contained in the hidden half of the H-maze, the proportion of trials in which participants revealed the hidden half of the maze was strongly correlated with their counterfactual processing noise indexed by performance in experiment 2 (r^2 = 0.80, p = 0.039, n=5). In contrast, no such correlation was found on trials in which the original half-maze included the correct exit point. These results reveal the bounded rationality of participants when relying on counterfactuals (Fig. 3c, right).

### Replication in large scale online experiments

The onsite experimental dataset was collected in a well-controlled lab setting but with limited participants (N=16). To validate the robustness of results, we pre-registered our hypotheses and analyses at the Open Science Framework (*OSF*, 2024) and repeated the experiments on an online platform (Prolific) using a larger pool (N=150). In online experiments, we were unable to measure eye movements and reliable reaction time data due to hardware limitations.

We first analyzed the online participants’ choice patterns on the baseline H-maze. Similar to onsite experiments, the counterfactual strategy best captured all participants’ choice pattern (Fig. 4a, similar to Fig. 2d, Left-Tailed T-test, p<1e-6 for all comparisons, n=49). Next, we analyzed online data from the three additional experiments to examine cognitive constraints. In *Task Variant 1,* participants were unable to process multiple streams of evidence in parallel effectively (Fig.4b, similar to Fig. 3a, Left-Tailed T-Test: p<2.67e-17, n=42). In *Task Variant 2*, similar to the onsite experiments, the estimated noise was higher when the maze was presented after the cue (Fig. 4c, similar to Fig. 3b, two-tailed T-Test: p = 7.4e-3, n=43). Finally in *Task Variant 3*, the proportion of trials in which participants elected to reveal the hidden half of the maze was strongly correlated with their counterfactual processing noise but only when the correct exit point was not in the hidden half (Fig. 4d, similar to Fig. 3c, Red: r^2 = 0.175, p = 5.2e-3, Blue: r^2 = 1.0e-4, p =0.96, n=43). In sum, the online experiments with the larger subject pool replicated the results from onsite experiments.

**Figure 4.**
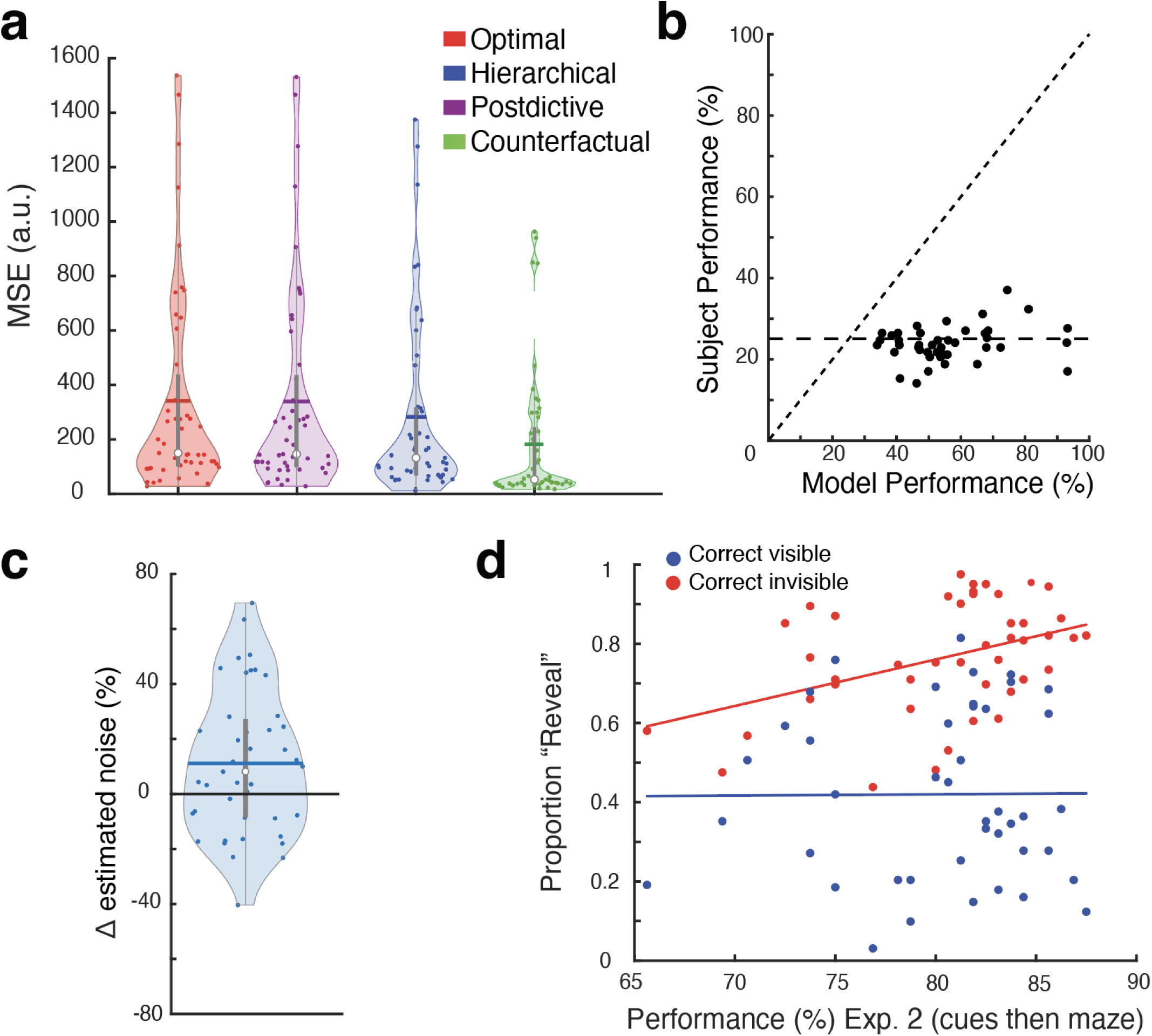
Large-scale online psychophysical experiments. **(a)** H-maze. Violin plots of the mean squared error computed between models’ and participants’ choice probability cross-validated five-fold. Colored circles are the mean MSE for each participant, colored solid horizontal lines are the mean MSE, the white circles indicate the median MSE, and the solid vertical gray bar indicates the 25^th^ and 75^th^ percentile MSEs. The MSE of the counterfactual model was significantly lower than other models in capturing participants’ choice patterns (Left-Tailed T-test, p<1e-6 for all comparisons, n=49). **(b)** Task Variant 1. Performance for individual participants compared to an ideal observer model solving experiment 1 using fitted *w* values measured separately for each participant on a control task (horizontal black dashed line: chance-level performance; diagonal black dashed line: unity). Performance of subjects was below the level expected by an ideal observer (Left-Tailed T-Test: p<2.67e-17, n=42). **(c)** Task Variant 2. Violin plot of the percent change in the estimated noise between condition 2 and condition 1. The estimated noise significantly increases in condition 2 compared to condition 1 (Blue circles: individual subjects, colored solid horizontal line: mean percent change in estimated noise, white circle: median percent change in estimated noise, solid vertical gray bar: the 25^th^ and 75^th^ percentiles, two-tailed T-Test: p = 7.4e-3, n=43). **(d)** Task Variant 3. The proportion of trials that participants revealed the hidden side of the maze as a function of performance in Condition 2 of Experiment 2 (Cues then maze) across participants. Blue circles are trials where the correct answer was on the revealed half of the H-maze, and red circles are trials where the correct answer was on the unrevealed half of the H-maze. The solid red and blue lines are the best linear fit to the data (Blue: r^2 = 1.0e-4, p =0.96, Red: r^2 = 0.175, p = 5.2e-3, n=43).

### A neural network model of counterfactual behavior

Behavioral responses in H-maze task variants revealed three characteristics of human decision-making: 1) an attentional bottleneck that limits processing multiple streams of information in parallel, 2) counterfactual processing noise that causes suboptimal performance, and 3) a judicious use of counterfactuals that takes this processing noise into account. Next, we used a modeling approach to ask whether these constraints provide an explanation for the participants’ cognitive strategy. Specifically, we asked whether imposing these constraints on recurrent neural network (RNN) models optimized to perform the H-maze task would lead to behavioral response patterns that are compatible with humans.

Architecturally, the model receives two types of inputs, has 128 recurrent units, and generates two outputs (Fig. 5a). The first type of input is a 6D vector with values associated with the geometry of the H-maze (*L_in_*, *R_in_*, *LU_in_*, *LD_in_*, *RU_in_*, and *RD_in_*). The second type of input, *I_time_*, is a 2D vector specifying the time intervals demarcated by the three auditory cues (*tm_1_*, *tm_2_*). The first output specified the model’s horizontal choice (*L_out_* versus *R_out_*), and the second output, the vertical choice (*U_out_* versus *D_out_*) (Fig. 5a).

**Figure 5.**
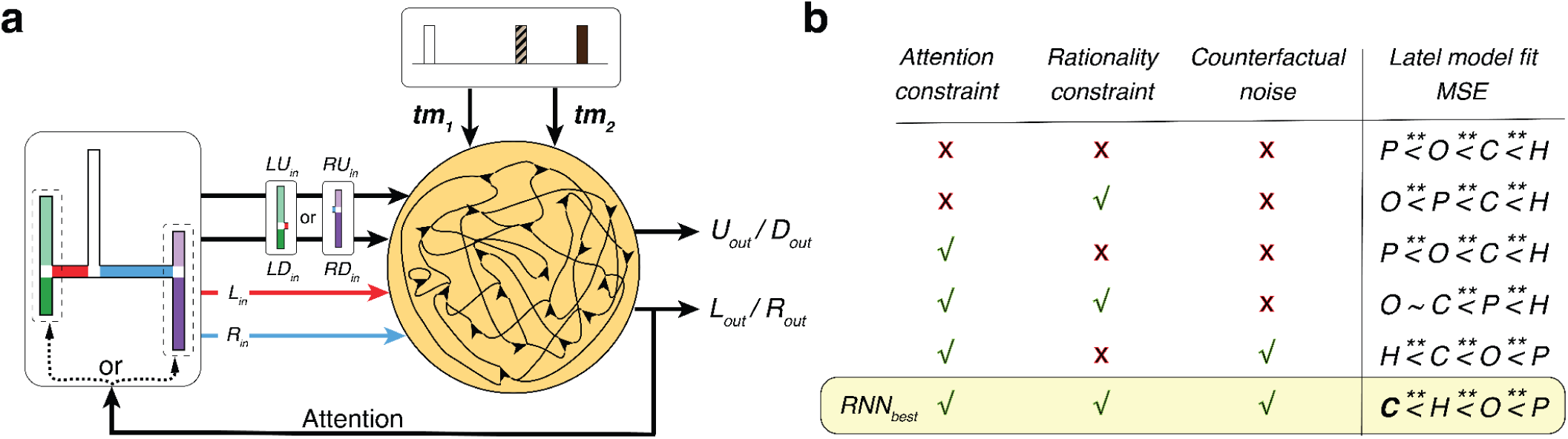
RNN with different constraints implement different decision strategies. **(a)**. RNN architecture. The RNN receives two types of inputs, a 6D vector specifying the arm lengths of the H-maze (*L_in_*, *R_in_*, *LU_in_*, *LD_in_*, *RU_in_*, and *RD_in_*) and a 2D vector specifying the noisy timing information for interval between the consecutive auditory cues (*tm*_1_ and *tm*_2_). The RNN produces two outputs. The first output specified the model’s horizontal choice (*L_out_* versus *R_out_*), and the second output, the vertical choice (*U_out_* versus *D_out_*). The horizontal output also serves as an attentional bottleneck forcing the RNN to choose between the vertical arms on one side of the maze, which changes the input it receives. **(b)** MSE computed on the choice probabilities between RNN variants and four latent models using a state-space analysis (see Methods). We trained variants of the RNN using different combinations of attention-bottleneck, rationality and counterfactual noise constraints. The table shows a subset of RNNs trained with (‘✓’) or without (‘✘’) certain constraints. Then for each of the RNN variants, we compared its choice pattern with the four latent behavioral models and ranked the models by their similarity to the RNN. Asterisks(**) indicate ‘significantly less than’ (p<0.05, one-tailed t-test, statistics calculated on 50 simulations). The yellow band highlights the RNN with all three constraints present, which is also the only RNN variant fitted best exclusively by the counterfactual model. We refer to this variant as RNN*_best_*.

Using this architecture, we developed various task-optimized RNNs subject to different sets of constraints. The base model was trained to choose the correct exit without any additional constraints. In the absence of any timing noise, this model could solve the H-maze perfectly without any errors. However, a noise-free model is unrealistic, as it does not account for the scalar variability that influences human timing behavior. We thus altered *I_time_* by adding scalar noise to *tm_1_* and *tm_2_* with Weber fraction, *w*=0.15, which is within the range observed across our participants (Fig. 2c, inset). As expected, introducing timing noise caused the model to make errors. In addition to the base model, we developed various task-optimized RNNs subject to one or more computational constraints inspired by the three experiments on participants’ decision-making strategy (Fig. 3). The constraints we considered were:

1. Attentional bottleneck: This constraint modifies training such that the RNN first makes a left/right choice (*L_out_* versus *R_out_*) and then uses this choice to *attend* to either the left or the right pair of vertical arms. We implement this constraint using a soft attentional gate that biases the input associated with the vertical arms to the left or right side of the H-maze (see Methods). In effect, this constraint forces the RNNs to solve the task hierarchically.
2. Counterfactual processing noise: This constraint modifies training such that the timing information the RNN relies on becomes progressively less reliable for each counterfactual revision. We implement this constraint by adding noise to *tm_1_*and *tm_2_* upon each revision (see Methods). This modification is analogous to assuming that the system is subject to counterfactual processing noise.
3. Rationality: This constraint modifies training such that the RNN learns to choose the exit that is most consistent with its noisy time interval measurements (as opposed to the correct exit). We implement this constraint using a cost function that enforces a maximum-likelihood decision strategy (see Methods). This is, in effect, a self-consistency constraint because decisions are made only based on information that is available to the model. Note that the base model violated this rationality constraint since it was optimized with perfect labels that were not available to the learner.

We compared the pattern of responses of each RNN to the pattern expected from the counterfactual model that best captured the behavior of human participants using a state-space analysis (Fig. 5b, see Methods). The response pattern of the base RNN was most dissimilar to humans. Next was the RNN with the rationality constraint, which fared better than the base RNN but was still quite dissimilar to the counterfactual model. Third was the RNN that was subjected to both the rationality and the attention-bottleneck constraints. Finally, the RNN whose response pattern was most similar to humans was the one that was subjected to all three constraints. We will refer to this RNN subjected to all constraints as RNN*_best_*.

Next, we took a closer look at the RNN*_best_* to assess its behavior more carefully (Fig. 6). We first verified that the overall performance of RNN*_best_*was similar to human participants and that its responses were sensitive to both horizontal and vertical arm-length differences (Fig. 6a). Next, we analyzed the “attention” signal in RNN*_best_* within single trials to assess whether the network relied on a counterfactual strategy. A purely hierarchical strategy would predict that RNN*_best_* would choose left or right after *tm_1_*and one of the corresponding vertical arms after *tm_2_*. In other words, under the hierarchical strategy, RNN*_best_* would never switch its attention signal to the alternate horizontal arm after *tm_2_*. In contrast, under the counterfactual strategy, RNN*_best_* would occasionally switch its left/right attention depending on arm lengths and *tm_2_*.

**Figure 6.**
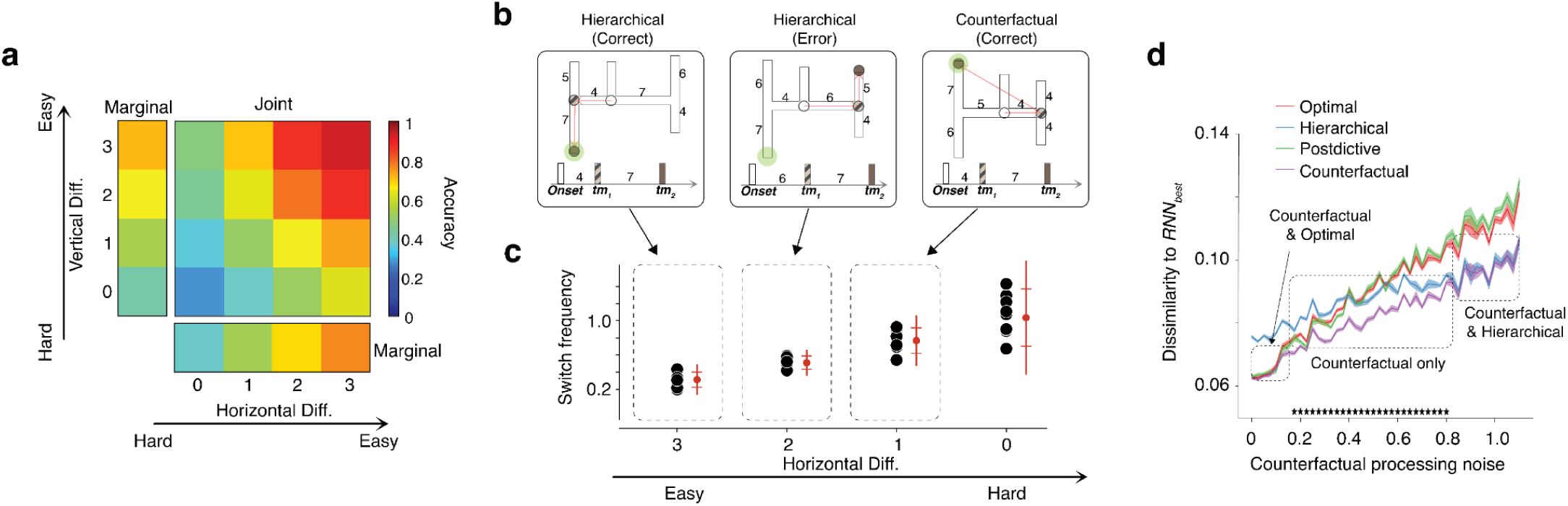
The counterfactual network (*RNN_best_*). **(a)** Behavioral performance. The accuracy of the *RNN_best_* improved systematically as a function of the difference between competing horizontal and vertical arms. **(b)** Single-trial behavior of *RNN_best_* for three geometries associated with different arm lengths and different measured time intervals (*tm*_1_ and *tm*_2_). Left: an easy trial where *tm*_1_ and *tm*_2_ match the left bottom exit. Middle: *tm*_1_ and *tm*_2_ are inconsistent with all exits; the RNN first chooses rightward based on *tm*_1_ and then upward based on *tm*_2_. Right: *tm*_1_ and *tm*_2_ are inconsistent with all exits; the RNN first chooses rightward based on *tm*_1_, but upon seeing *tm*_2_, switches its attention to left and chooses the left-up as its final choice. The two red arrows trace *RNN_best_* decisions after *tm*_1_ and *tm*_2_. The green circle shows the correct answer. See Methods for details. **(c)** The probability of attentional switches as a function of the first choice difficulty (i.e., the difference between the horizontal arms is smaller). Results are sampled from 10 networks. The three trials in **(b)** are samples of the three horizontal differences highlighted in **(c)** (guided by arrows). **(d)** Dissimilarity between the response pattern of *RNN_best_* and various cognitive models as a function of counterfactual processing noise (σ*_noise_*). The shading shows the standard deviation across 50 simulations. Asterisks indicate the conditions in which *RNN_best_* is most similar to the counterfactual model. The dashed blocks show regions with low, intermediate and high σ*_noise_*, which specify the cognitive model to which *RNN_best_*is most similar to.

We found that RNN*_best_* single trial behavior was indeed consistent with the counterfactual strategy. When the difference between horizontal arm lengths was large, RNN*_best_* adopted a hierarchical strategy without any revisions (Fig.6b, left and middle). In these conditions, the final decision was usually correct when the difference between the corresponding vertical arms was large (Fig.6b, left), and sometimes wrong when the vertical arms were more similar (Fig.6b, left). Remarkably, when the difference between the horizontal arm lengths was smaller, RNN*_best_*would sometimes revise its first left/right decision (as inferred from the attention signal) depending on *tm_2_* (Fig.6b, right). This attention-switching behavior increased systematically for smaller differences between the horizontal arms for which the first decision was more uncertain (Fig.6c). These behavioral characteristics provide direct evidence that RNN*_best_* solves the task using a combination of hierarchical and counterfactual strategies.

Finally, we probed the parametric influence of counterfactual processing noise on the behavior of RNN*_best_*. Similar to our Experiment 3 in humans, (Fig. 3c), we predicted that the frequency with which RNN*_best_* would rely on counterfactuals would decrease for higher levels of counterfactual processing noise. Results were consistent with this prediction; the proportion of attentional switches dropped systematically under higher levels of counterfactual processing noise. One insight gleaned from this analysis was that the network’s inferred strategy was qualitatively different for small and large amounts of counterfactual processing noise (σ*_noise_*). On one end of the spectrum, when noise levels were small (σ*_noise_*<0.18), the network could rely on counterfactuals without any cost, and thus its behavior was indistinguishable from the optimal model (Fig. 6d). On the other end of the spectrum, when noise levels made counterfactual processing too costly (σ*_noise_*>0.8), the network rarely relied on counterfactuals, and thus its behavior was indistinguishable from the hierarchical model. In other words, computations RNN*_best_* were adaptive and exhibited a graceful transition from optimal to counterfactual to hierarchical with the counterfactual strategy emerging as the best solution for intermediate levels of noise (0.18<σ*_noise_*<0.8; p<0.05, one-tailed t-test comparing counterfactual to the next best model).

Together, these results indicate that human participants’ strategy is consistent with the predictions of the bounded rationality hypothesis governed by an attentional-bottleneck and the magnitude of counterfactual processing noise. Moreover, the continuous transition between strategies implies that the distinct cognitive algorithms could be mere subdivisions in a strategy continuum under the same objective function.

## Discussion

Humans’ cognitive capacity limitations make them unable to find optimal solutions when facing moderately complex problems. Yet, we are quite efficient at finding reasonably good solutions. In cognitive sciences, this ability is viewed through the lens of computationally rational bounded optimality, which posits that humans rely on strategies that are optimal within the bounds of their computational capacity. When facing large decision-trees, one of the most common strategies humans use is to think through a hierarchy of if-then scenarios and, when needed, consider counterfactuals. While numerous studies have found evidence for these strategies (Huys et al., 2015; Lake et al., 2017; Tenenbaum et al., 2011; Zylberberg, 2021), less is known about the computational constraints that motivate these strategies. Here, we used a moderately complex task for which we could generate precise models to predict behavior under optimal, bounded-optimal and suboptimal strategies. Comparing model predictions to behavior, we were able to verify that humans used a combination of hierarchical and counterfactual strategies.

Next, we performed a series of complementary experiments to gain insight into the potential constraints that underlie these strategies. In a first experiment, we verified that parallel processing degrades performance, consistent with previous findings (Davis & Marcus, 2016; Ludwin-Peery et al., 2021; Smith et al., n.d.). This result is consistent with previous work showing humans’ inability to process multiple streams of information simultaneously, and the importance of an attentional bottleneck to address this limitation (Dasgupta et al., 2020; Duncan, 2013; Gershman & Burke, 2022; Ramírez-Ruiz & Moreno-Bote, 2022; Wolfe et al., 2006; Zylberberg, 2021). In a second experiment, we quantified performance degradation due to counterfactual processing, which is likely due to capacity limitations of memory-based computations (Beck et al., 2012; Egger & Jazayeri, 2018; Panichello et al., 2019; Ratcliff, 1978; Remington et al., 2018; Sarafyazd & Jazayeri, 2019; Shadlen & Shohamy, 2016; Yu et al., 2020; Zylberberg, 2021). In a third experiment, we made the intriguing observations of lower likelihood of counterfactual reasoning for subjects for whom counterfactuals incurred a larger performance cost. This finding suggests that human’s use of counterfactuals is not optimal but computationally rational (Gershman et al., 2015; Piloto et al., 2022). Together, the results of these experiments provided a clear hypothesis for when humans rely on hierarchical and counterfactual processing: hierarchical processing may be used to compensate for performance degradation due to parallel processing and counterfactual processing may be used judiciously when it can improve performance.

However, human experiments alone do not provide a strong test of these hypotheses because we cannot experimentally manipulate cognitive capacity limitations and computational constraints in participants. To address this shortcoming, we developed task-optimized RNNs that were subjected to different computational constraints, and analyzed their behavioral patterns to see whether there might be a direct link between computational constraints and cognitive strategies. This modeling effort proved highly fruitful. It provided clear evidence that the models that were subjected to constraints identified in humans found solutions to the H-Maze task that were consistent with hierarchical and counterfactual processing, similar to humans. Moreover, the parametric sweeps of the model revealed a direct link between working memory limitation and counterfactual processing. Finally, we found that a single model with a fixed architecture may generate behavior consistent with optimal strategy, hierarchical strategy, or counterfactual strategy, depending on the computational constraints and noise-levels. It is therefore conceivable that strategies that are typically considered as distinct within cognitive sciences may be part of a continuum within neural systems (Donoso et al., 2014). This finding may open new avenues of exchange between adjacent studies of the brain in systems neuroscience and the mind in cognitive sciences.

## Materials and Methods

### Human participants

Fifteen participants (age: 18 to 65 y, six male and five female) participated in the experiments after giving informed consent. All participants were naive to the purpose of the study, had normal or corrected-to-normal vision, and were paid for their participation. All experiments were approved by the Committee on the Use of Humans as Experimental Subjects at the Massachusetts Institute of Technology.

Five participants completed the baseline H-Maze task, five participants completed task variant experiment one, and five participants completed task variant experiments two & three. The testing sequence for the two tasks was counterbalanced across participants. In each session, a participant was seated in a dark quiet room and asked to perform the task of interest for ∼60 min. For both tasks, stimuli and behavioral contingencies were controlled by an open-source software (MWorks; mworks-project.org/) running on an Apple Macintosh platform.

### Extended online participants

In the extended online version of the task, we recruited 150 more participants (age: 18 to 70 years old, 87 males and 63 females) on Prolific (www.prolific.com). The participants gave consents and the protocol was approved by the Committee on the Use of Humans as Experimental Subjects at the Massachusetts Institute of Technology.

The participants were divided into three groups with 50 participants in each group. Participants in group 1 performed the T-maze (168 trials) and H-maze (540 trials) tasks sequentially in one session (∼60 minutes). Participants in group 2 performed the T-maze (168 trials) and Task Variant 1 (170 trials) sequentially in one session (∼30 minutes). Participants in group 3 performed the Task Variant 2 (320 trials) and Variant 3 (324 trials) sequentially in one session (∼60 minutes). For Task Variant 3, the hidden side of the maze was always on the right side, but the correct answer could be either on the left or right side. The tasks were coded in jsPsych (www.jspsych.org) and deployed on the Cognition.run (www.cognition.run) platform.

#### Exclusion criteria

To ensure the quality of online participant data, we exclude subjects whose measured Weber fraction *w* on the control T-maze experiment or condition 1 of experiment 2 exceeds 0.4.

### Tasks

#### Baseline experiment and naming conventions

The main stimulus in the baseline experiment was a ball moving inside a maze shown from above at a constant speed. The ball entered the maze from the top through a vertical entry hallway, continued moving through the maze at a constant speed, and stopped at one of many possible exit points. In the majority of experiments, the maze was shaped like the letter H, hence the name H-maze. In the H-maze experiment, the vertical entry hallway branched into 2 horizontal arms each of which branched into 2 vertical arms. We will use these subscripts to refer to the length of specific arms (*a_L_*, *a_R_*, *a_LU_*, *a_LD_*, *a_RU_*, *a_RD_*) as well as the time it would take the ball to move along those arms (*t_L_*, *t_R_*, *t_LU_*, *t_LD_*, *t_RU_*, *t_RD_*). The initial hallways was 7 degree visual angle (dva), and the H-maze arms lengths were sampled from a discrete uniform distribution with values 3, 4, 5, 6, and 7 dva subject to the constraint that the sum of all arm pairs (L+R, LU+LD, RU+RD) was 10 dva (bottom left). Accordingly, the absolute difference between each arm pair (|L - R|, |LU - LD|, |RU - RD|) could be either 0 (5, 5), 2 (6, 4), or 4 (7, 3) dva (right). The speed of the ball was 8 or 12 dva/sec. With these parameters, the shortest and longest arms lengths (3 and 7 dva) were associated with 0.25 and 0.875 sec, respectively. The width of all arms was 0.5 dva.

In the H-maze, the ball (1) started moving downward along the entry hallway until reaching the horizontal segment, (2) turned left or right and continued at the same speed until reaching one of the vertical segments, (3) turned up or down and continued at the same speed until reaching one exit point where it stopped. Throughout the manuscript, we use subscripts *L* (left) and *R* (right) to refer to the left and right arms of the horizontal segment. Moreover, we will use subscripts *LU* (left-up), *LD* (left-down), *RU* (right-up), and *RD* (right-down) to refer to the 4 corresponding vertical arms.

#### H-maze experiment

The trial structure in the H-maze experiment was as follows: (1) An H-maze was presented for 1.5 s. Together with the maze, four circles marking the four exit points were presented (color: white; diameter = 1 deg) (2) a ball visibly traveled through the initial hallway at one of two randomly sampled speeds. (3) Three identical auditory cues were presented (cue duration: 80ms). The first cue was presented when the ball reached the horizontal segment, the second cue when the ball reached the corresponding vertical segment, and the third cue when the ball reached the end of the maze. We will refer to the interval between the first and second cues as the first sample interval, denoted by *ts_1_*, and the interval between the second and third cues as the second sample interval, denoted by *ts_2_*. (4) Participants had to make a keyboard choice (q for LU, a for LD, p for RU, and l for RD) to indicate the exit point corresponding to the end position of the ball. The trial would abort if no answer was made within 2500 ms. (5) The circle corresponding to the participant’s choice changed color to provide feedback (green if correct, purple if incorrect). (6) Trials were separated by an inter-trial interval of 1.5 sec.

#### T-maze experiment

The T-maze experiment is identical to the H-maze experiment except that there are vertical arms and the exit points are at the end of the two horizontal arms (*L* and *R*). Accordingly, participants had to report their binary decision about the exit by choosing one of two circles presented at the end of the two horizontal arms. The two arm lengths (*a_L_*, *a_R_*) were sampled from the same distribution as the H-maze. The speed of the ball was also the same as the H-maze task. Therefore the distribution of the time intervals associated with the two arms (*t_L_*, *t_R_*) was also the same as the H-maze task. We performed this experiment to estimate each participant’s timing variability parameterized by a weber fraction, *w* (see below).

#### Task variant 1

Task variant 1 was designed to force participants to implement the optimal model, where the participant updates their beliefs about all four possible ball trajectories simultaneously. To do so, (i) participants were presented with a variant of H-maze where the four possible ball pathways are flattened into four parallel options, and the timing statistics (e.g. total time of occluded ball movement) were kept identical between the H-maze task and task variant 1 ; (ii) after a 1.5 second viewing period, four balls initialized in the same location moved visibly toward the maze with identical and constant speed; (iii) an auditory cue signaled when the balls reached the horizontal arms at which time the balls were made invisible and continued to move rightward. If a ball reaches the left or right end of its arm, the ball reverses direction. Crucially, the 1) nonlinearity introduced by the ball direction reversal dissociated the balls’ position from time and thus forcing participants to separately keep track of four ball trajectories separately, and 2) no cue was provided at the time of any ball direction reversal, thus participants cannot solve this task hierarchically or sequentially; (iv) a second auditory cue signaled when one the balls reached the endpoint of its arm. Crucially, the arm segments and the number of direction reversals were chosen so that the three other balls were not at any arm segment endpoint and were still traveling through their respective arm segments. The endpoint of the ball could be on the right or left end of the arm and after a variable number of direction reversals; (v) Participants had to click one of four circles that appeared next to four possible end points of the ball, and participants had to indicate at which arm end pint (vi) participants received a binary visual feedback at the location of choice.

#### Task variant 2

Task variant 2 is similar to the H-maze experiment except that only one half of the maze is shown to the participants (*a_L_*, *a_LU_*, *a_LD_* or *a_R,_ a_RU_*, *a_RD_*), and thus there are only two exit points similar to the T-maze experiment. The occluded ball traveled through the horizontal arm shown and one of the two vertical arms shown. Accordingly, participants had to report their binary decision about the exit by choosing one of two circles presented at the end the vertical arms. All arm lengths were sampled from the same distribution as the H-maze. The speed of the ball was also the same as the H-maze task. Therefore the distribution of the time intervals associated with the two arms (*t_L_*, *t_R_*) was also the same as the H-maze task. In condition one, the participants were shown the half maze and then the auditory temporal cues related to ball movements, just as in the baseline task. In condition two however, the order of stimulus presentations were reversed: the half maze was presented after the auditory temporal cues related to ball movements, forcing participants to maintain sensory evidence in working memory and use it afterwards to test different hypotheses about the subsequently shown half maze (i.e., counterfactual reasoning). All arm lengths were sampled from the same distribution as the H-maze. The speed of the ball was also the same as the H-maze task.

#### Task variant 3

Task variant 3 is similar to the task variant 2 where initially only one half of the maze is shown to the participants (*a_L_*, *a_LU_*, *a_LD_* or *a_R,_ a_RU_*, *a_RD_*), however the true generative process of the ball’s movements were identical to the baseline H-maze task (the ball could travel through the revealed horizontal arm and one of the revealed vertical arms, or through the hidden horizontal arm and one of the two hidden vertical arms). The participants were shown the half maze and then the auditory temporal cues related to ball movements. Participants decided the exit point among the two available options in the half maze, and subsequently were given the option to reveal the hidden half of the maze, which would allow them to revise their decision and choose between the two new exit ports (i.e., revision via counterfactual reasoning). All arm lengths were sampled from the same distribution as the H-maze. The speed of the ball was also the same as the H-maze task.

#### Decision model

Every time the ball reached a bifurcation point, it took one of two alternative arms. We developed a simple decision model based on signal detection theory (Green & Swets, 1966) to characterize how participants discriminated between the two arms using the time between the corresponding two cues. Let us denote the two arm lengths by *a_1_* and *a_2_*. Since the ball moves at a constant speed, the arm lengths correspond to the two time intervals, *t_1_* and *t_2_*. Depending on which arm the ball takes, the interval between the two cues, denoted *t_s_*, will either match *t_1_* or *t_2_*. The participant has to use a noisy measure of the actual interval, denoted *t_m_*, to decide which of two arms the ball has taken.

Consistent with scalar variability of timing (Malapani & Fairhurst, 2002), we modeled noise in the measurement of time as a sample from a zero-mean Gaussian distribution whose standard deviation scales with the base interval (*t_1_*or *t_2_*), with the constant of proportionality *w*. According to this model, the conditional probability of *t_m_* for the two arms can be written as:

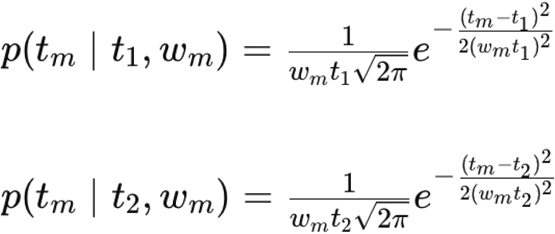

Using signal detection theory (Green & Swets, 1966), the optimal decision is to choose the arm with the higher conditional probability. This is equivalent to choosing the side on which *t_m_* falls relative to a criterion, *t_c_*, at the crossing point of the two conditional probability distributions. We refer to this model as the core decision module.

#### Estimating each participant’s Weber fraction

We estimated *w* for each participant by fitting the core decision module to the behavior in the T-maze task. To accurately estimate *w*, we included *t_c_* as a model parameter to account for idiosyncratic biases. The model also included an extra lapse rate parameter, to account for participants’ lapses in performance. During lapse trials, the model chooses randomly between the two arms. With these additional parameters, the conditional probability for each arm can be written as follows:

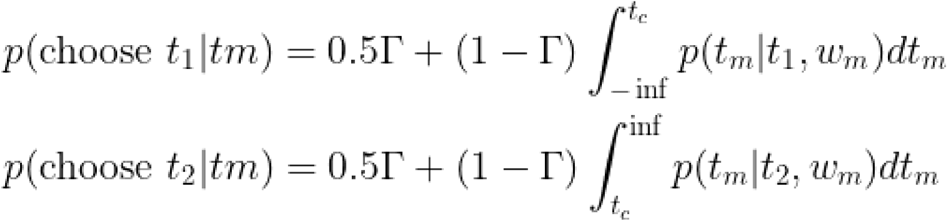

We computed the maximum likelihood estimate of *w_m_* (as well as the other two parameters) for each participant by fitting this model to behavior during the T-maze task.

#### Optimal model

This model chooses the most likely exit by comparing the joint conditional distribution of *tm_1_* and *tm_2_*for the four alternatives. The four conditional probabilities are can be written as *p(tm_1_,tm_2_|t_L_,t_LD_)*, *p(tm_1_,tm_2_|t_L_,t_LU_)*, *p(tm_1_,tm_2_|t_R_,t_RU_)*, and *p(tm_1_,tm_2_|t_R_,t_RD_)*.

We computed these joint conditional probabilities by the product of corresponding marginals assuming that the two measurements are conditionally independent. Note that this model has no additional parameters other than *w* that was estimated from the T-maze experiment. Therefore, we did not fit this model to the participants’ behavior; instead, we evaluate its behavior predictively.

Hierarchical Model.

The hierarchical model solves the H-maze task by making two decisions, first between the horizontal arms and then between the corresponding vertical arms. The first decision chooses between the two horizontal arms by comparing the corresponding conditional probabilities, *p(tm_1_|t_L_)* and *p(tm_1_|t_R_)*. The model chooses the more likely arm and then uses *tm_2_* to choose between the two vertical arms that branch off of the chosen horizontal arm. For example, if the model chooses the left arm in the first stage, it would then compare *p(tm_2_|t_LD_)* and *p(tm_2_|t_LU_)* to choose between the left-up and left-down vertical arms. This model also has no additional parameter other than *w*, which was estimated from the T-maze experiment. Therefore, the behavior of this model was also evaluated predictively.

#### Postdictive Model

The postdictive model is similar to the hierarchical model in that it solves the task by making two hierarchical decisions, first for the horizontal arms and then for the corresponding vertical arms. However, in this model, the first decision for the two horizontal arms is made postdictively by incorporating additional information about the *tm_2_* and the vertical arms. Specifically, the model makes its first decision by comparing *p(tm_1_,tm_2_|t_L_,t_LD_)*+*p(tm_1_,tm_2_|t_L_,t_LU_)* to *p(tm_1_,tm_2_|t_R_,t_RU_)*+*p(tm_1_,tm_2_|t_R_,t_RD_)*. Next, it uses *tm_2_* to choose between the two vertical arms that branch off of the chosen horizontal arm, which is identical to the second decision in the hierarchical model. Similar to the optimal and hierarchical model, this model has no additional parameter (*w_m_* is derived from the T-maze experiment).

#### Counterfactual Model

The counterfactual model is identical to the hierarchical model but has the flexibility to revise its decisions when a certain measure of expected accuracy (*X*) is lower than a certain threshold (*θ*) that was fit as a free parameter.

#### Noise in the counterfactual model

To compute the likelihood of a counterfactual possibility, participants must perform an offline mental computation that involves inferring the values of *tm_1_* and *tm_2_* from memory to test a new hypothesis (the likelihood that the exit is down the other horizontal arm). Since the process of inferring time intervals from memory is noisy, we formulated the counterfactual model such that the recalled values of *tm_1_* and *tm_2_* (values used during the revision process) were participant to additional noise. Similar to the noisy optimal, hierarchical, and postdictive models, the noise in the counterfactual model was sampled from a Gaussian distribution with mean *β* and standard deviation that is proportional to the measured interval (*tm_1_* or *tm_2_*) with a fixed constant of proportionality *α* (in accordance with scalar variability of time). The parameters *α* and *β* were fitted using maximum likelihood estimation.

#### Choice state-space analysis

For the state-space analysis, we encoded the pattern of choices in terms of four probabilities, one for each possible exit, denoted by *p_LU_*, *p_LD_*, *p_RU_*, and *p_RD_*. We first applied isometric transformations (rotation and reflection) to all H-mazes on each trial such that the left-up exit was the correct answer for all trials (the correct exit point of the ball was always *LU)*. *p_LU_* is the probability of the correct exit point which is the most likely option (for analysis the H part of the maze on each trial was reflected horizontally and/or vertically so that the correct exit point of the ball was always *LU*), *p_LD_* is the probability of choosing the correct horizontal arm but incorrect vertical arm, and *p_RU_* and *p_RD_* are the probabilities of choosing the incorrect horizontal and vertical arms. Next, we grouped unique trial conditions into a smaller number of maze sets. Each maze set contained all the mazes with the same arm length differences. We then quantified the dissimilarity between the behavior of each model and the participants/RNNs for each maze set by measuring the distance between the corresponding 3D vectors composed of [*p_LU_ p_LD_ p_RU +_ p_RD_*]. Finally, we averaged the distance across maze sets to compute an overall measure of dissimilarity between choice patterns of the participants/RNNs and each model. To account for the limited number of trials from human subjects, we grouped trials into a smaller set of maze conditions based on only the horizontal arm time difference for our human analyses.

#### Estimated noise

In our pre-registration, we proposed a model-free analysis of experiment 2 data based on subject performance for different arm length differences. However, this analysis confounds performance differences due to stimulus difficulty (arm length difference) with the timing sensitivity of subjects. Here we conduct an improved model-based analysis of experiment 2 data where we fit a performance-based psychometric curve over all arm length differences for each subject. Let us denote the two arm lengths by *t_1_* and *t_2_,* and the difference in measured time between *t_1_* and *t_2_ as t_d_*. To characterize how participants discriminated between the two arms using the time between the corresponding two cues, we fit a cumulative gaussian model to compute the probability of a correct choice given *t_1_*and *t_2_* as follows:

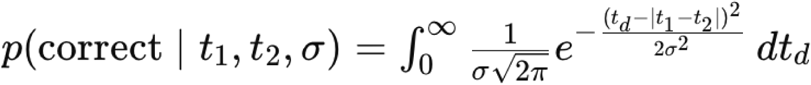

Once we compute the maximum likelihood estimate of each subject’s noise σ for each condition, we calculate the percent change in the subjects’ estimated noise σ between condition 1 and condition 2 as:

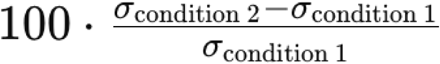

The hypothesis that subjects’ estimated noise increases on condition 2 relative to condition 1 predicts a significantly positive percentage change. The null hypothesis predicts no such significant change.

#### The RNN model

To construct the trials for the RNN, we sample the length of each of the six maze segments [*L, R, LU, LD, RU, RD*] from the set of lengths {4, 5, 6, 7} uniformly. Thus there are 4^6 = 4,096 possible maze geometries. We randomly split the geometries into 3,072 (75%) for model training and 1,024 (25%) for model testing. When RNN performs the task, we apply noise to the inter-click intervals. We convert each ground truth interclick interval *ts_i_* to a perceived inter-click interval *tm_i_* = *ts_i_* · (1 + σ*_noise_*), where ε ∼ N (0, 1) for σ*_noise_* = 0.15. This noise is independently sampled for each of the two inter-click intervals per stimulus. For each maze geometry, we consider each of the four possible ball paths and for each of the conditions sample 10 perceived inter-click interval pairs (*tm*_1_, *tm*_2_). Thus we generate 40 trials for each maze geometry in both the training and testing sets.

To train RNNs on the H-maze task, we discretize time into 32 timesteps per trial. We let the ball begin 5 length units above the horizontal maze arms and travel at 1 unit per timestep, so on every trial the ball reaches the first T-junction at timestep 6. For auditory input, we construct a horizontal and vertical auditory timing variable [*a*_1,_ *a*_2_]. We let *a*_1_ be 0 until the ball reaches the first T-junction, then ramp linearly from 0 to the first perceived inter-click interval *tm*_1_ in *int*(*tm*_1_) (discretize to nearest integer) timesteps and remain constant thereafter. Similarly, we let *a*_2_ be 0 until *a*_1_ reaches *tm*_1_, upon which *a*_2_ ramps linearly to the second perceived inter-click interval *tm*_2_ and in *int(tm_2_)* timesteps then remains constant. For visual input, we provide models with scalar values of the arm lengths [*L, R, LU, LD, RU, RD*]. The RNN provides two outputs, a horizontal choice and a vertical choice ([*h, v*]) that jointly specify the chosen exit.

Our counterfactual RNN model consists of a recurrent state ***b*** of *N* = 128 hidden units, updated each time step according to

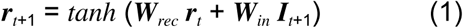

where ***W**_rec_* is an *N* × *N* matrix of trainable recurrent weights, ***W**_in_* is a *N* × 8 matrix of trainable input weights, and ***I*** is a 8-dimensional input vector [*a_1_*_*t*_, *a_2_*_*t*_, *L, R, LU, LD, RU, RD*] at each timestep.

The model produces two choices (horizontal and vertical) at each timestep representing the vertical and horizontal choices:

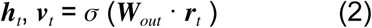

where ***W**_out_* is an *N* × 2 matrix of learnable weights and σ is the sigmoid function σ (σ) = 1/(1 + *e*^−*x*^). We use a threshold of 0.5 to binarize the choice to evaluate behavioral outcomes.

For the basic version of the RNN, we use an external-supervising signal to train the RNN. For each trial, we minimize the MSE between the RNN output [*h_t_*, *v_t_*] and the ground truth output. The loss is summed over the timesteps after *tm*_2_ finishes ramping.

#### Attention constraint

To add the attentional bottleneck to the RNN, we change the original 8-dimensional input [*a_1_*_*t*_, *a_2_*_*t*_, *L, R, LU, LD, RU, RD*] into a 6-dimensional input [*a_1t_*, *a_2t_*, *L,, R, U_*t*_, D_*t*_*], where *U_t_* and *D_t_* are the lengths of vertical arms gated by an attention module. Mechanistically, the attention module produces outputs *U_t_* and *D_t_* with the horizontal choice ℎ_*t*_ at each timestep *t* as follows:

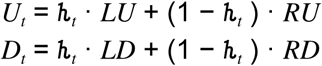

Accordingly, the input weight matrix becomes *N* × 6 with the attention constraint.

#### Rationality constraint

Instead of supervised learning, we train the RNN with the objective of maximizing the likelihood of the chosen exit given the noisy timing information, which is the rational choice under uncertainty. Given a maze geometry and perceived inter-click intervals *tm*_1_ and *tm*_2_, an rational policy on the H-maze task chooses an exit-point corresponding to a path in the maze that maximizes the joint log likelihood log (*p* (*tm*_1_|*t*❘_1_)) + log (*p* (*tm*_2_|*t*❘_2_)). Because *tm_i_* is sampled from a Gaussian distribution centered at the ground truth *ts_i_*, maximizing the log likelihood is equivalent to minimizing the loss function

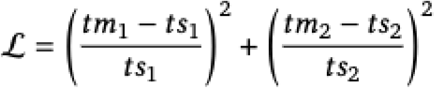

Considering the output logits [***h**_t_*, ***v**_t_*] as the model’s choice of which side of the maze the ball exits (left/right and up/down), the loss function becomes

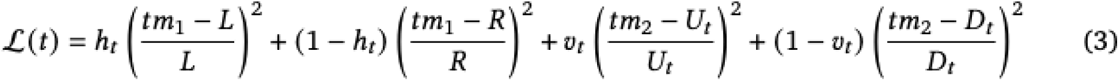

On the right-hand-side of the equation, the first two terms are the loss coming from the horizontal arm choice *h_t_* and the last two terms are the loss coming from the vertical arm choice *v_t_* given the horizontal arm choice. When training the models, this loss function is summed over the timesteps after *tm*_2_ finishes ramping.

#### Counterfactual noise constraint

In order to discourage the model from switching its attention from side to side every trial, we impose an implicit cost for switching: after the second auditory click (namely after *a_2_*finishes ramping), each time ℎ crosses 0.5 we add noise to the perceived inter-click intervals [*tm*_1_, *tm*_2_]. Specifically, for each timestep *t* where ℎ crosses 0.5, we update *tm* as

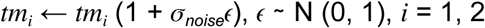

and use this noised *tm* for all subsequent timesteps in the trial. Here σ*_noise_*is a hyperparameter controlling the magnitude of this switching penalty. We set σ*_noise_* = 0.5 for the counterfactual noise constraint and σ*_noise_* = 0 without this constraint.

### RNN variants and model comparison

(Fig. 4b). There are in total eight possible RNN variants with three constraints that are binary (attention, rationality, and counterfactual noise). However, we ruled out two illogical variants: since counterfactual noise is triggered by the reverse of attention, it does not make sense to have counterfactual noise without the attention constraint. When comparing each RNN variant with the counterfactual model, we need a maximum likelihood estimation to find the best noise level of the counterfactual model. For simplicity, we simulated the counterfactual model three times with different noise levels: σ*_noise_*=0 (no noise), σ*_noise_*=0.5 (medium noise), σ*_noise_*=1 (high noise). We chose the noise level for the counterfactual model that produces the least MSE to the RNN.

### RNN saccade

(Fig. 5b). Although the RNN generates continuous outputs [*h_t_*, *v_t_*] throughout the trial, we plot the saccade at two discrete time points for clarity (the time steps when *tm*_1_ and *tm*_2_ finishes ramping). For the first saccade (right after *tm*_1_), we only consider *h_t_* and ignore *v_t_* because the RNN cannot make a vertical decision without *tm*_2_. For the second saccade (right after *tm*_2_), the saccade position reflects both *h_t_* and *v_t_*.

## Author contributions

M.R. and M.J. conceived the study. M.R. collected the human psychophysics data and performed all the analyses. C.T., N.W., M.R. and M.J. conceived the RNN architecture. C.T. and N.W. implemented the RNN code. C.T. collected the online experiment data. C.T. performed all RNN analyses. M.R., C.T., N.W. and M.J. wrote the manuscript. M.J. supervised the project.

## Data availability

The data used to generate the associated figures will be made available on a public repository after peer-review publication.

## Code availability

The code used to generate the associated figures will be made available on a public repository after peer-review publication.

## Acknowledgments

M.R. is supported by Lisa K. Yang ICoN fellowship. C.T is supported by Friends of the McGovern Institute Student Fellowship. N. W. is supported by the National Science Foundation Graduate Research Fellowship Program. M.J. is supported by the Simons Foundation, the McKnight Foundation, and the McGovern Institute.

## Supplementary Figures

**Supplementary Figure 1.**
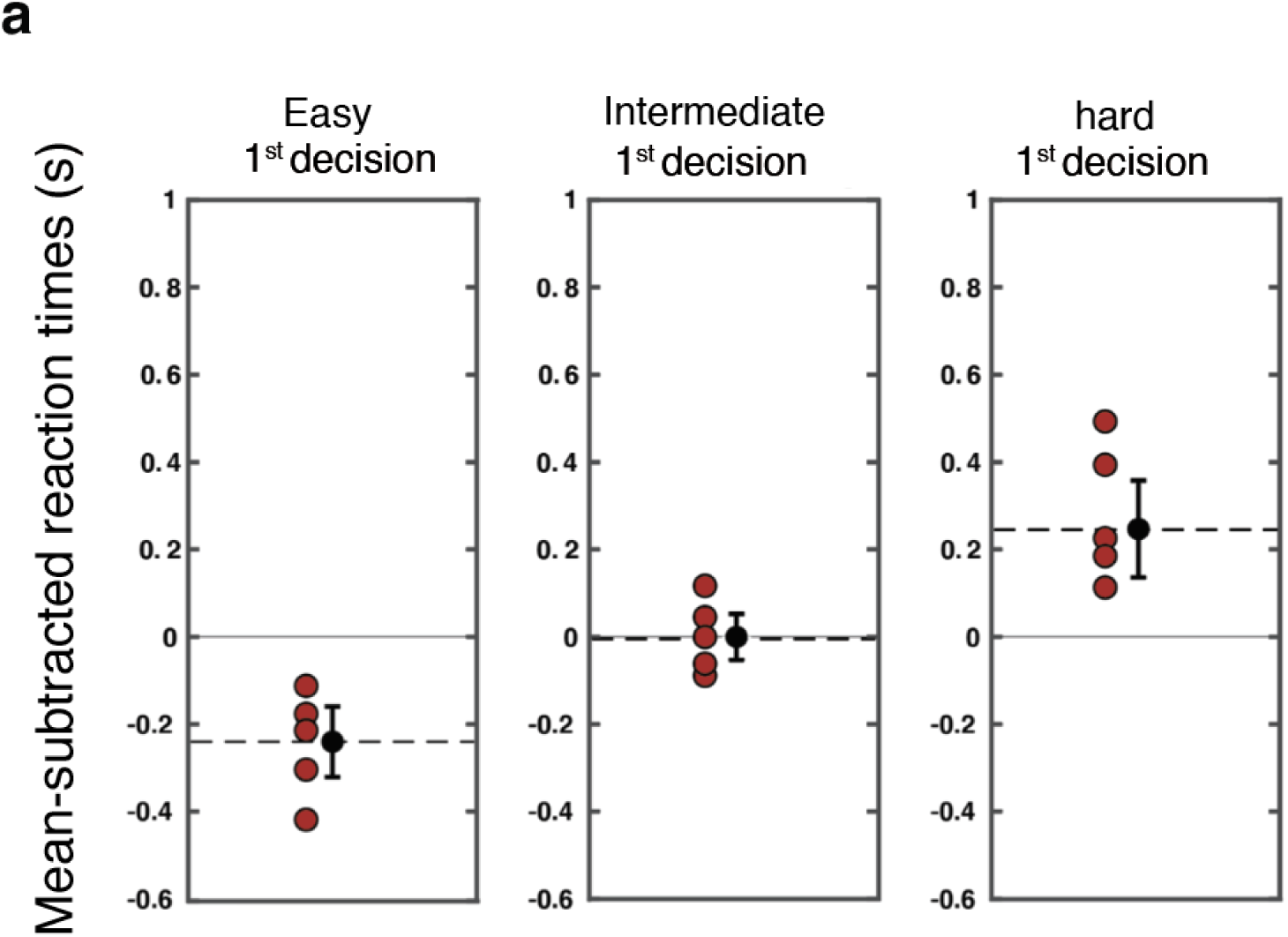
Participants’ reaction times on Baseline H-Maze. **(a)** Mean subtracted reaction times (computed as the time from flash three to the subject’s choice) of human participants (seconds) for 1st decision conditions with low, intermediate, and high difficulty are plotted in red. Average reaction times of participants per condition is plotted in black, with std. indicated by black error bars.

**Supplementary Figure 2.**
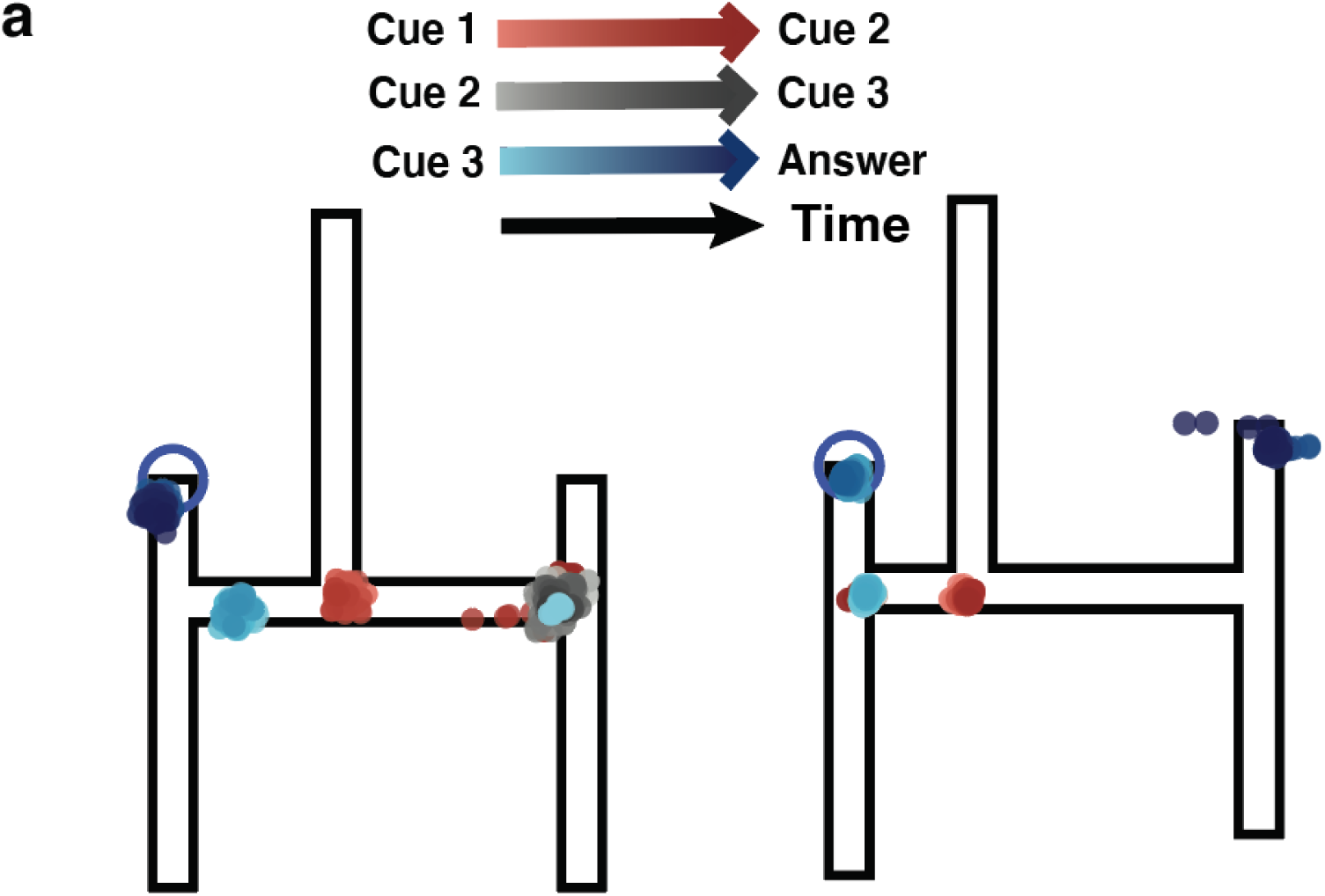
Dynamic Eye-Movement Readout. **(a)** Gaze position density throughout the trial is overlaid on the H-maze. In these examples, the correct endpoint is top-left and the participant’s answer is indicated by the open blue circle. Gaze position density is color coded based on the task epoch (red: the ball is in the horizontal segment; gray: the ball is in the vertical segment; blue: the trial has ended and the ball has reached the endpoint). Bright and dark shades represent early and late in each epoch. The pattern of eye movements indicate that participants made their choice hierarchically (early leftward or rightward saccades in the top left panel). They changed their mind occasionally and made a saccade to the other vertical bar either after the second auditory cue (top right), or the third auditory cue (bottom left and right).

**Supplementary Figure 3.**
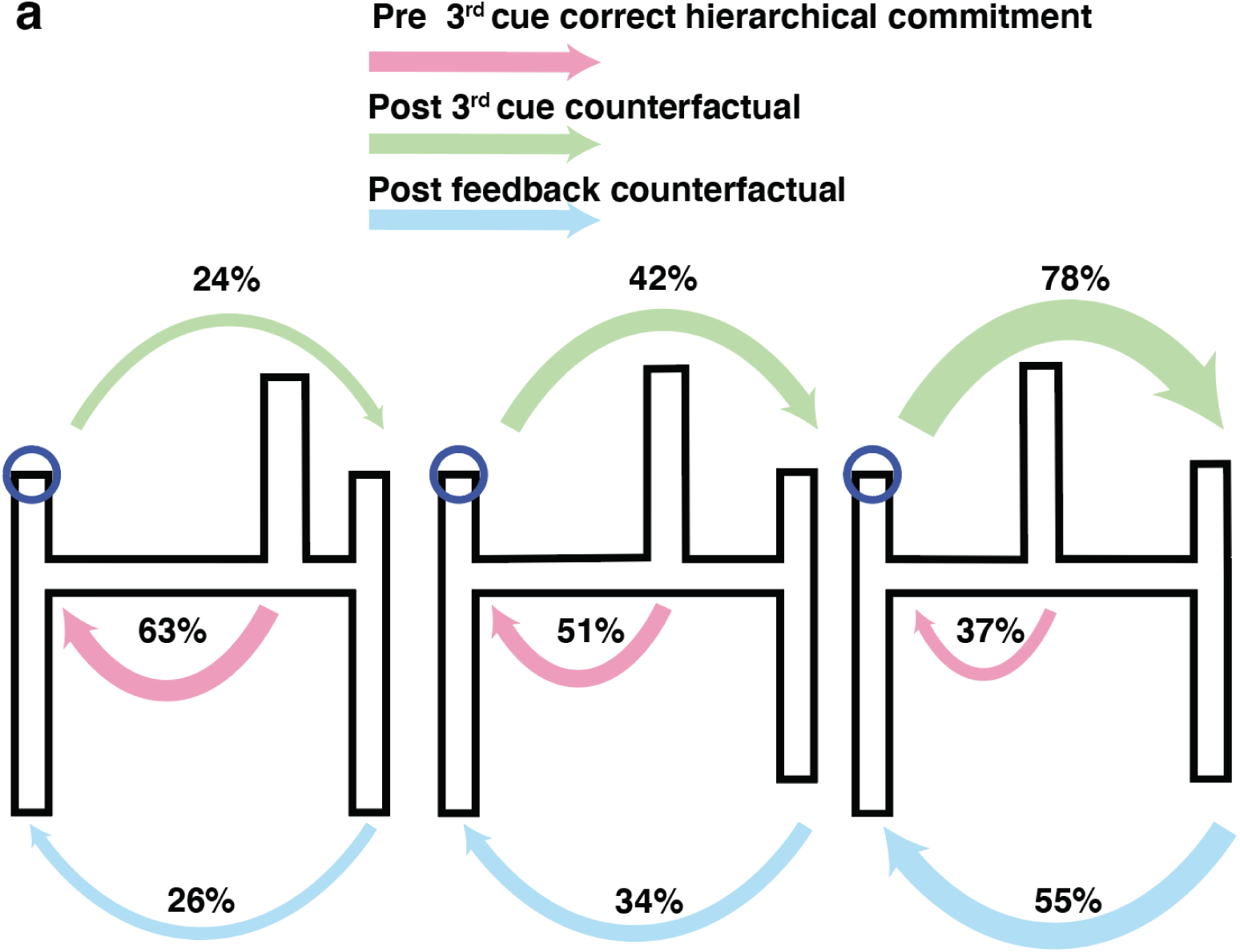
Condition dependent eye-movements. **(a)** A directed probabilistic graph of the transition probabilities between hierarchical and counterfactual saccades is shown for the three horizontal arm difficulty conditions, plotted separately here on three maze schematics representing each condition. The pink arrows represent correct hierarchical saccades made to the one of the two horizontal arms before the third cue is presented. The green arrow represents counterfactual saccades made after the third cue was provided to the arms of the maze alternative to the initial decision commitment. The blue arrow represents saccades to the arms made after choice feedback to the arms of the maze alternative to the initial decision commitment. The width of arrows is proportional to probability, and the percentage of trials on which the transition occurred is shown.

