## Supplementary figures and images for "Computational basis of hierarchical and counterfactual information processing"

### Supplemental Figure 1

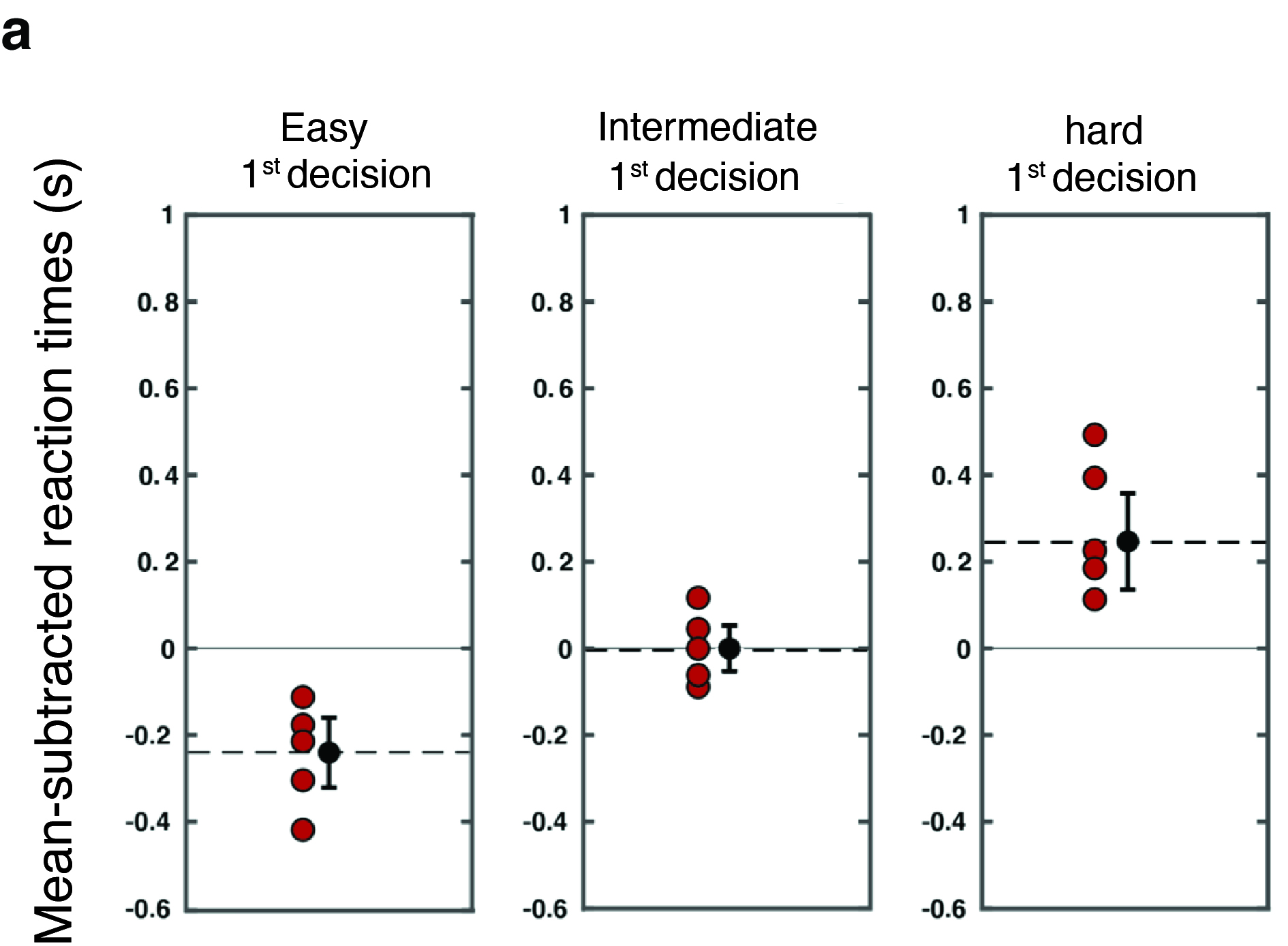

### Supplemental Figure 2

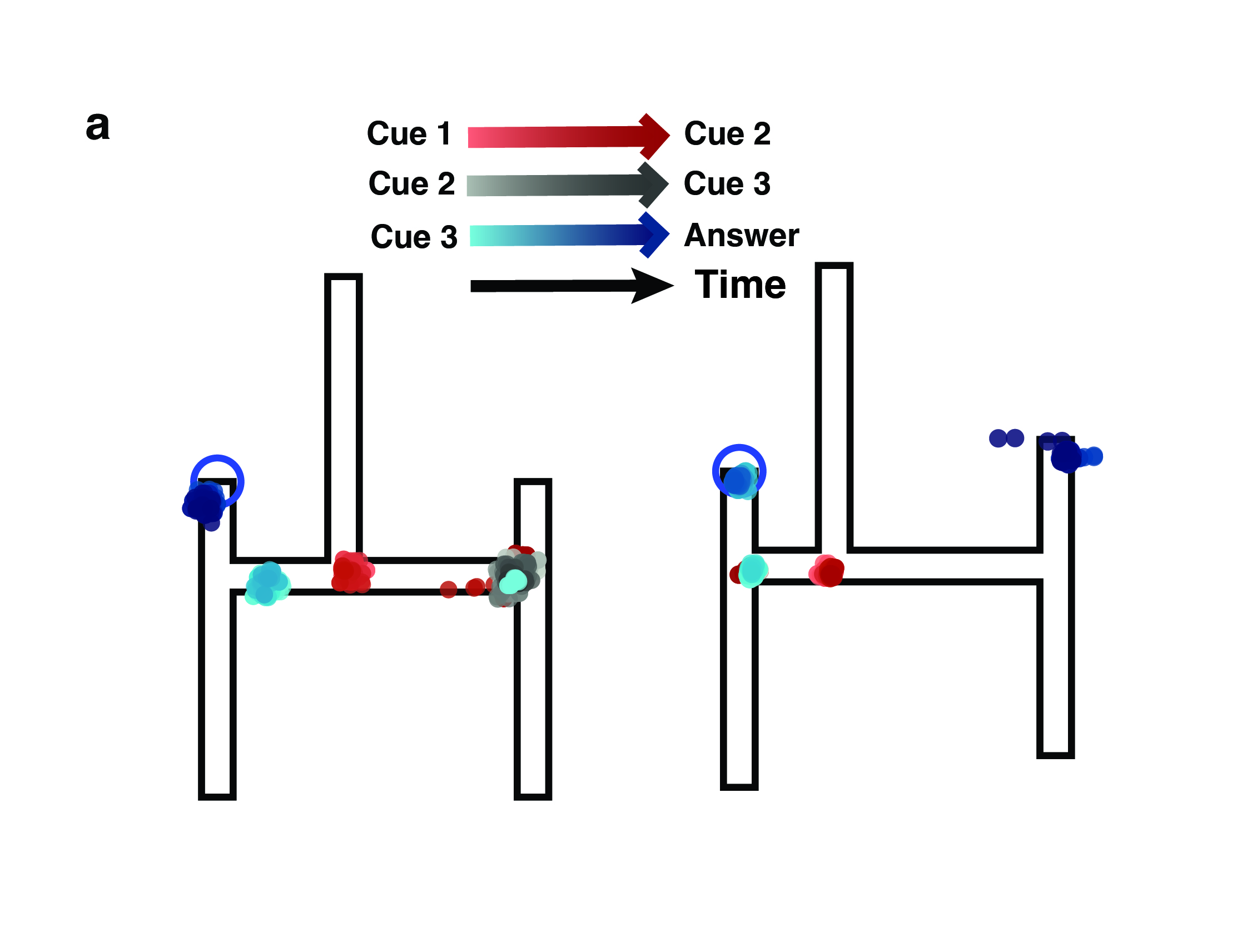

### Supplemental Figure 3

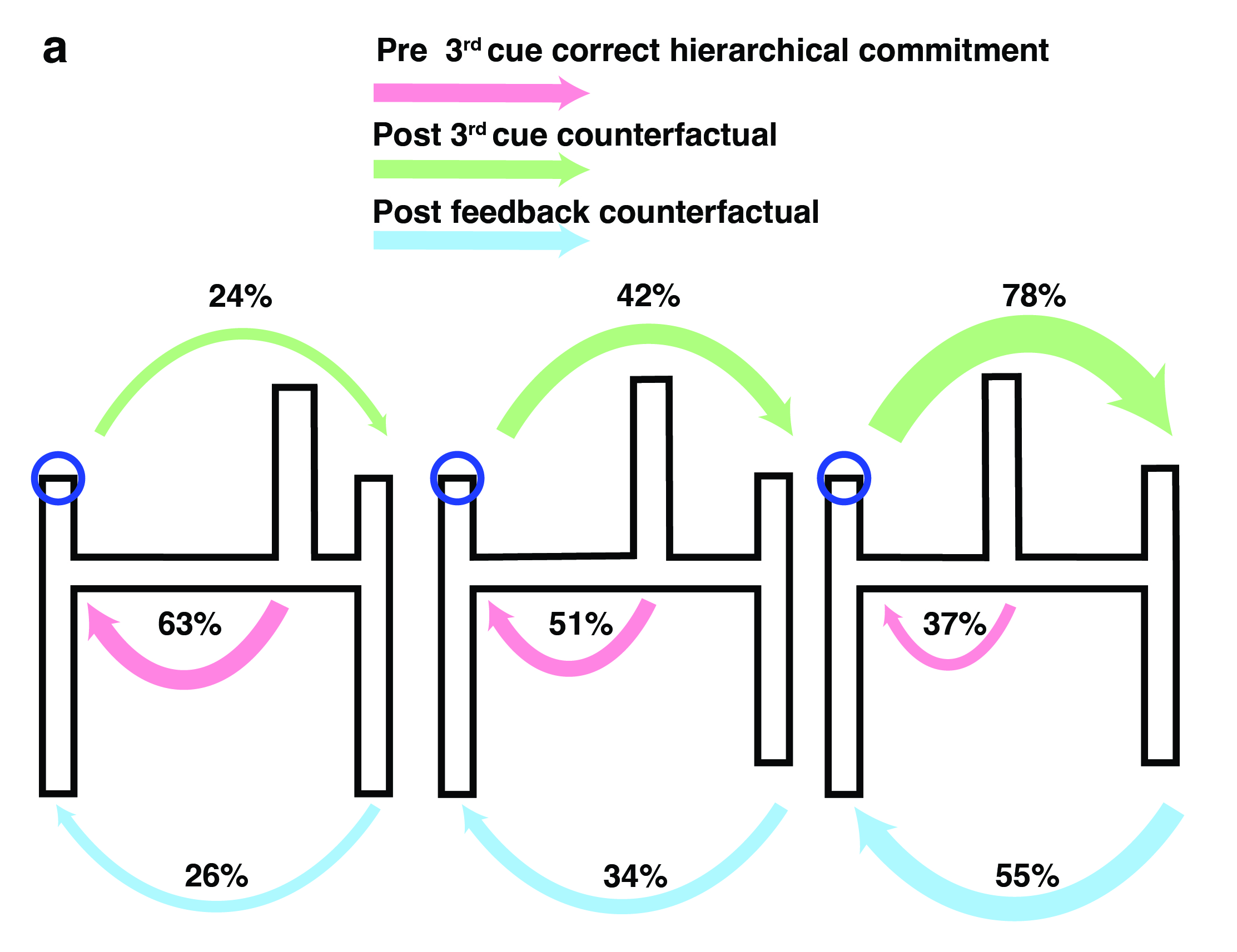
